# FAST-STEM: A Human pluripotent stem cell engineering toolkit for rapid design-build-test-learn development of human cell-based therapeutic devices

**DOI:** 10.1101/2024.05.23.595541

**Authors:** Aaron H. Rosenstein, Rangarajan Sambathkumar, Brandon M. Murareanu, Navroop K. Dhaliwal, Fumao Sun, Xinyaun Zhao, Abolfazl Dadvar, Rasha Al-attar, Andrew Chai, Nitya Gulati, Ting Yin, Maria Nguyen, Danielle Serra, Tania Devina, Aanshi Gandhi, Mohammad Saleh, Penney Gilbert, Tilo Kunath, Michael A. Laflamme, Shinichiro Ogawa, Julien Muffat, Yun Li, Stephanie Protze, Cristina Nostro, Michael Garton

## Abstract

Very recent clinical advances in stem cell derived tissue replacement and gene therapy, in addition to the rise of artificial intelligence-aided scientific discovery, have placed the possibility of sophisticated human cell-based therapies firmly within reach. However, development of such cells and testing of their engineered gene circuit components, has proven highly challenging, due to the need for generating stable cell lines for each design–build–test–learn engineering cycle. Current approaches to generating stable human induced pluripotent stem cell (hiPSC) lines are highly time-consuming and suffer from lack of control, poor integration efficiency, and limited functionality. Validation in clinically relevant stem cell derived tissues is also broadly lacking. Such drawbacks are prohibitive to repeatably conducting cutting-edge stem cell engineering with broad application within a realistic timeframe and will not scale with the future of regenerative medicine. We have developed FAST-STEM (***F****acile **A**ccelerated **S**tem-cell **T**ransgene integration with **S**ynBio **T**unable **E**ngineering **M**odes*), a hPSC engineering platform that drastically reduces the time to generate ‘differentiation ready’ stem cell lines from several weeks to 5 days, exhibiting a ~612-fold improvement in transgene integration rate over previous methodologies. Additional FAST-STEM innovations include: (i) rapid and highly efficient transgene integration; (ii) copy number control; (iii) simultaneous or consecutive integration of multiple gene cassettes; (iv) library screen capability. In addition to this unique functional versatility, platform transportability and broad use case for stem cell-engineering was confirmed by differentiation into eight different cell types across nine different laboratories. This platform dramatically lowers the bar for integration of synthetic biology with regenerative medicine, enabling experiments which were previously deemed logistically impossible, thus paving the way for sophisticated human cell device development.

## Introduction

Tissue replacement therapy using human pluripotent stem cell-derived products is on the cusp of becoming a clinical reality for treatment of numerous diseases such as heart failure, diabetes, multiple sclerosis, and Parkinson’s^1–4^. Beyond direct tissue replacement, this advancement raises the possibility of designing cells with novel therapeutic functionality that can be safely engrafted into human recipients. Example applications could be neuronal cells that detect and respond to seizures, or ‘factory’ cells that make and secrete therapeutic proteins in response to disease cues. Human cells are an ideal chassis for therapeutic device design due to their biocompatibility, versatility, and ability to interact dynamically with the human body. They can receive a wide variety of inputs, from chemical and biological ligands, to light, time, pressure, electrical stimulation, and more ^5–11^. Cells can be programmed to integrate these inputs to compute a huge variety of outputs and actuations, including molecule secretion, proliferation, differentiation, chemotaxis, event recording and more ^12–15^. Using this ever-growing mammalian synthetic biology toolkit, incredibly compact and powerful ‘biological computers’ can potentially be programmed to achieve novel therapeutic effects^16–19^. Very recent advances in genetic component and genome design using generative machine intelligence ^20–29^, offer potential for even more sophisticated ideas, such as the creation of novel human cell types. However, despite their potential, prototyping with clinically relevant cells, in particular human pluripotent stem cells (hPSC) has been severely hampered by several major barriers. These include the immunogenicity of synthetic components, unpredictable context effects, resource overload, and epigenetic silencing^30–33^. An overarching challenge–– preventing timely solution development on all these fronts––is the paucity of engineering tools for hPSCs and the hitherto unavoidable slow pace of design iteration.

Both in induced human stem cells (hiPSCs) and human embryonic (hESCs), site specific, stable cell line generation typically takes a minimum of three weeks, with additional time spent on differentiation, typically ranging from one to six weeks and in some cases even longer^34,35^. Lentiviral, and transposase-based tools such as *piggyBac,* are relatively quick and efficient means of gene integration in hPSCs, producing stable populations ready for differentiation in ~10 days. However, integration occurs at random genomic loci with variable efficiency, leading to unpredictable levels of transgene expression, silencing, and impact on endogenous genetic elements, often yielding unrepeatable results^36,37^. Positional effects from random integration events have been shown to result in transgene silencing in hiPSCs^38–40^. The alternative, CRISPR-Cas9, is a highly targeted approach that can produce stable transgene integration via homology-directed repair (HDR) at defined genomic loci in hPSCs, obviating many of these issues. However, it generally takes months or more to produce, validate, and bank differentiation ready cells, and the screening workload necessary can be prohibitively time-consuming and costly. CRISPR knock in can also be cytotoxic and exhibit reduced efficiency in comparison to target-agnostic methods like *piggyBac*^37,41,42^.

To address these challenges, hPSCs have been engineered with ‘landing-pads’; exogenously-derived recombination sites at loci with predictable activity and minimal silencing ^43–48^. Trans-genesis using a delivery vector is equal or more efficient than direct CRISPR-Cas9-based transgene knock-in, less prone to unintended mutagenesis, less toxic, and can provide predictable stable transgene expression. Safe harbour sites such as *AAVS1, ROSA26, CLYBL*, and *SHS231* enable long-term stable transgene expression and are common landing-pad locations ^43,46,49–51^. Various mechanisms for effecting gene transfer into landing-pad cells have been characterized, leveraging Cre, Flp, ϕC31, and Bxb1-based recombination machinery^43–48,52^. More advanced methods utilize recombinase-mediated cassette exchange (RMCE), where parallel recombination events effect specific integration of only genetic material between two recombination sites, excluding integration of bacterial plasmid backbones^48^. Recombinase-based landing-pad targeting strategies in hiPSCs include Big-IN, STRAIGHT-IN, dual-integrase cassette exchange (DICE) system and ‘target attP’ (TARGATT™) ^43–45^. These have reduced the time taken to produce differentiation ready cells to approximately a month, although they lack validation of differentiation to clinically relevant tissues, and only offer limited knock-in options (single-gene) to the synthetic biologist/cell engineer.

In other organism contexts (bacteria, yeast, plant), many challenges in synthetic gene circuit development have been very effectively addressed by establishing standardized parts and performing rapid iterative design–build–test–learn (DBLT) cycles^53–55^. The same approach has also been effectively applied to human cells, where transient transfection and tractable immortalised cell lines such as HEK293 and HeLa have yielded many valuable tools^56^. However, circuits developed with such cell lines are often ineffective in hPSCs and derived tissues (e.g., transgene silencing of the cytomegalovirus promoter (CMV), which exhibits robust expression in many transformed cells^57–59^). DBLT cycles are highly resource intensive and time-consuming in hPSCs, as generation of stable cell lines is required for every iteration. Yet, such a capability is essential for effective integration of generative-AI-assisted genetic component design with regenerative medicine. Because of the large DNA or protein sequence spaces explored by many generative models (i.e. LLMs, VAEs, GANs) a novel system is required to deploy such experiments with single-cell resolution and readouts with deep-learning-scale magnitude.

Here, we present FAST-STEM (***F****acile **A**ccelerated **S**tem-cell **T**ransgene integration with **S**ynBio **T**unable **E**ngineering **M**odes*), an hPSC engineering platform that cuts the time to generate *differentiation ready* cell lines with repeatable transgene expression dynamics to 5 days. FAST-STEM further provides cell-based therapy engineers with the ability to simultaneously integrate two different transgenes into each AAVS1 locus and can receive combinatorial libraries for facile high throughput screening. FAST-STEM hPSCs were differentiated into eight different clinically relevant cell types across nine different laboratories, demonstrating transferability and broad applicability. Altogether, the speed, versatility, and reliability of FAST-STEM has the potential to catalyse the integration of generative AI and synthetic biology with regenerative medicine.

## Results

To generate a platform with the broadest possible utility to the regenerative medicine/cell-based therapy engineering community, we conducted a knowledge gathering exercise in these communities. Following this, iPS11 was chosen as the parental line, due to its ease of handling, and reliability in differentiation to many different tissue types of therapeutic interest. FAST-STEM constructs were designed and installed to this cell line as follows:

### FAST-STEM Construct Design

HDR-donor constructs were generated by assembly of either EGFP, tagBFP or mScarlet into an Ef1a-driven neomycin-resistance polycistron (Figure 1a). Constructs were flanked by Bxb1 *attB*(GT) and *attP*(GA) sites and homology arms for the *AAVS1* locus respectively. All constructs were verified by whole-plasmid sequencing to ensure functionality in downstream experiments.

**Figure 1:**
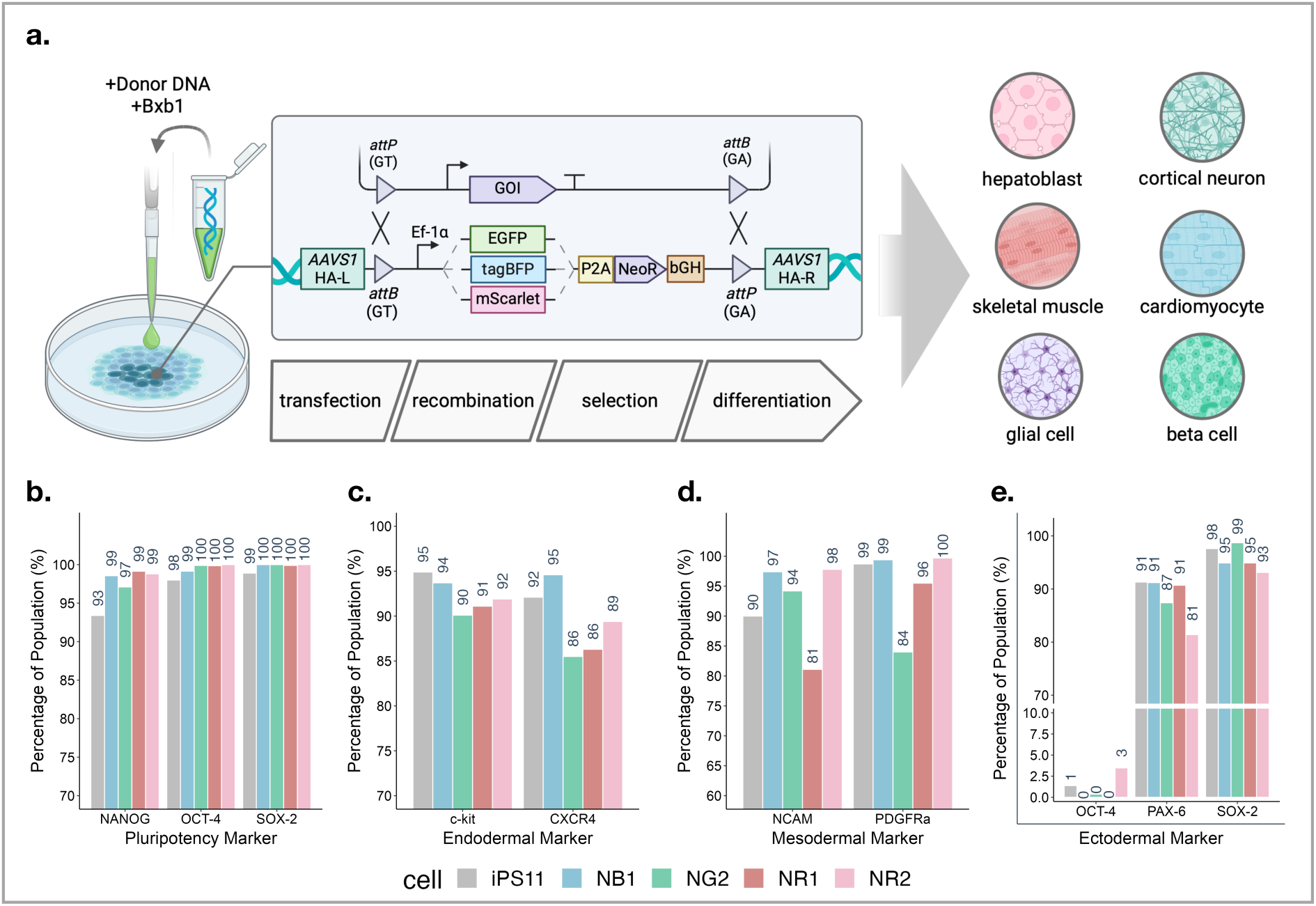
FAST-STEM construct design, pluripotency and differentiation marker profiling. **(a)** Diagram of FAST-STEM construct use, including transfection of hiPSCs with donor DNA and Bxb1, subsequent recombination into the genome with overall workflow, and tested differentiated tissue types described herein. A gene of interest flanked by compatible attP/B site pairs is depicted being targeted to the FAST-STEM. **(b)** Immunostaining and subsequent flow cytometric analysis of OCT-4, SOX-2, and NANOG expression markers compared FAST-STEM stemness to the parental iPS11 cell line. **(c)** Endoderm differentiation assessed by immunostaining of c-kit and CXCR4 expression. **(d)** Mesoderm differentiation assessed by NCAM and PDGFRa staining. **(e)** Ectoderm differentiation assessed by loss of OCT-4, gain of PAX-6, and maintenance of SOX-2 expression.

### CRISPR HDR-Mediated Construct Integration into *AAVS1 and* Verification Thereof

Delivery of FAST-STEM donor construct to hiPSCs followed by selection, clonal expansion, and fluorescence-activated cell sorting (FACS) yielded double-copy EGFP and mScarlet reporter cell lines (classified as NG2 and NR2 respectively) and single copy tagBFP and mScarlet reporter cell lines (classified as NB1 and NR1 respectively). Each were screened to ensure correct knock-in by PCR of the *AAVS1* locus with primers spanning regions distal to the repair template homology arms. In addition to droplet-digital polymerase chain reaction (ddPCR) of the Neomycin resistance gene (NeoR) and flow cytometric analysis (Supplementary Figure 1a, b), Nanopore sequencing of PCR products specific for the NG2, NR2, NB1, and NR1 FAST-STEM constructs followed by alignment with the CRISPR HDR repair-template plasmid confirmed correct insertion of all components. We noted varying degrees of loss of fluorescent protein expression following cell engineering and screening stages. As a test case, we conducted further FACS sorting of the NR2 population into mScarlet+/- populations, and tracked mScarlet expression over the course of 28 days in both conditions (Supplementary Figure 1c). Secondary copy-number analysis of the NeoR gene by ddPCR indicated that the FAST-STEM construct was maintained in both populations at a double-copy within the genome (Supplementary Figure 1d). These results indicate that transgene silencing may be the cause of a decline in reporter expression as opposed to construct loss from the genome.

### Maintenance of Pluripotency, Baseline Differentiation Potential, and Karyotype in FAST-STEM hiPSCs

To ensure that genetic manipulation of the *AAVS1* locus and monoclonal expansion did not impact FAST-STEM hiPSC pluripotency, verification of OCT-4, SOX-2 and NANOG expression was determined (Figure 1b). All FAST-STEM clones retained pluripotency marker expression in comparison to parental iPS11 cells. Differentiation of FAST-STEM cells into all three germ layers (endoderm, mesoderm ectoderm) was also assessed in comparison to parental iPS11. For endodermal marker profiling, immunostaining and flow cytometric analysis of CXCR4 and c-kit expression after four days of differentiation revealed all FAST-STEM cells exhibited double-positive populations similar to parental iPS11 cells (Figure 1c). Mesodermal profiling of NCAM and PDGFRα revealed all FAST-STEM cells exhibited similar staining profiles to iPS11 controls as well (Figure 1d). Ectodermal PAX-6, maintenance of SOX-2 and loss of OCT-4 profiling confirmed all FAST-STEM cells were also capable of directing differentiating to this lineage (Figure 1e). Furthermore, all FAST-STEM hiPSC lines retained reporter expression throughout differentiation processes, albeit with varying degrees of silencing (Supplementary Figure 1e). Karyotyping of the NG2, NR2, NB1 and NR1 FAST-STEM hiPSC lines revealed that cells were karyotypically normal following CRISPR-HDR and monoclonal expansion (Supplementary Figure 1f-i). Thus, these hiPSCs were deemed viable for testing of downstream applications.

### Developing a Flexible Repertoire of FAST-STEM Delivery Vectors

In order to easily deliver a variety of gene constructs into FAST-STEM hiPSCs, a standard collection of vectors with sequentially increasing degrees of complexity were created (Figure 2a). The most basic “level-0” vector only contained a multiple-cloning site (MCS) flanked by *attP*/*attB*’ recombination sites. The “level-1” vectors incorporate splice-acceptor-based antibiotic resistance (puromycin or blasticidin) or fluorescent protein (EGFP, tagBFP, mScarlet) gene-trap cassettes. Additionally, flanking the MCS with ALOXE-3 and cHS4 transgene insulator sequences yielded the complete set of level-2 vectors^60^. All vectors were designed for compatibility with the MTK toolkit, creating an interface with extant modular cloning workflows^61^.

**Figure 2:**
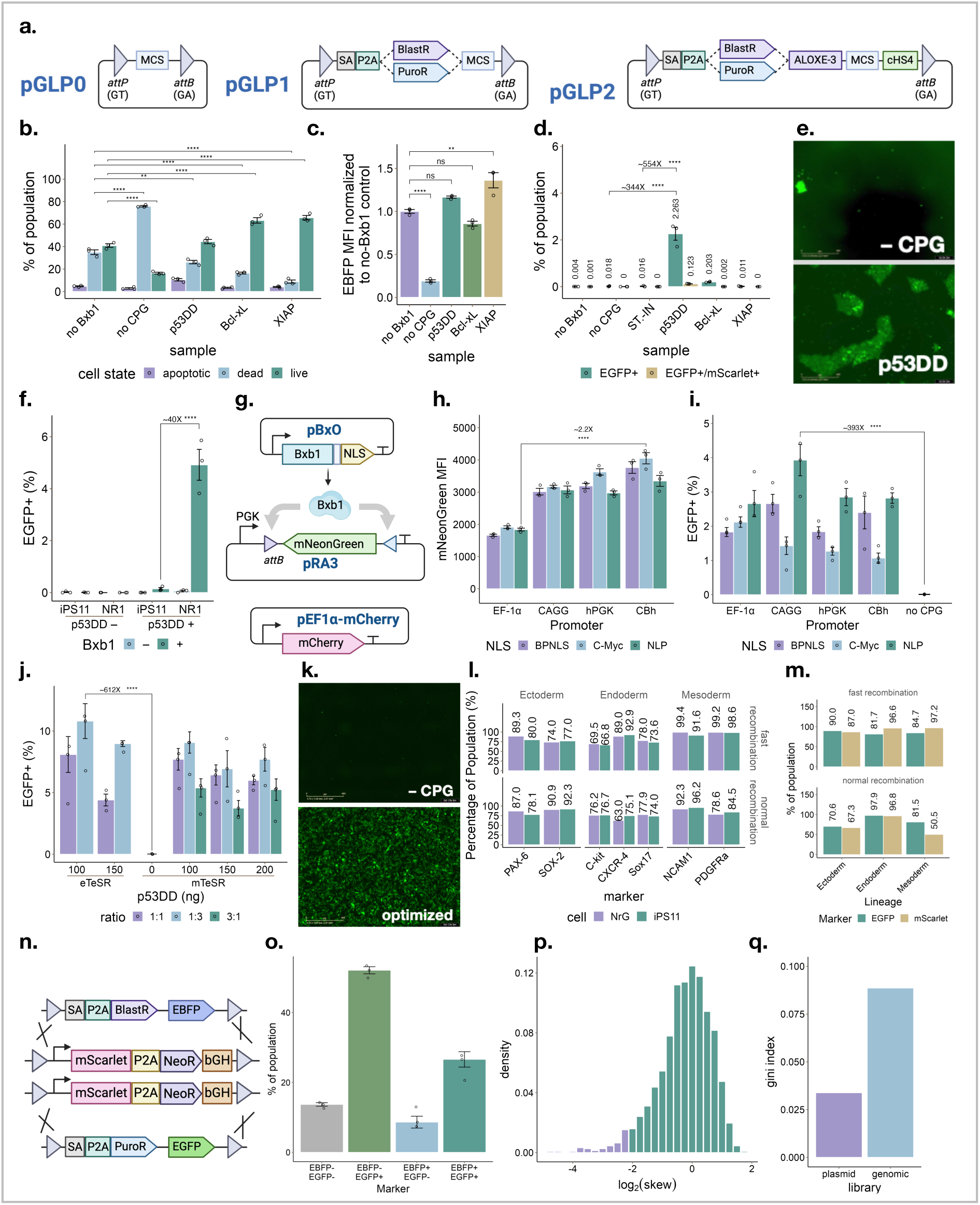
Enhancing transgene delivery to FAST-STEM hiPSCs and developing synthetic-biology-relevant engineering modes. **(a)** Schematic of delivery vector levels 0-3 are depicted with either puromycin and blasticidin antibiotic resistance markers. **(b)** Live/dead/apoptosis flow cytometric quantification as well as **(c)** transfection efficiency for NR1 cells transfected with an EBFP transgene with no Bxb1, no cytoprotective gene (CPG) or one of three CPGs (p53DD, Bcl-xL, XIAP). **(d)** Proportion of cells expressing EGFP transgene in NR1 cells 10-days post-transfection. **(e)** Representative fluorescence microscopy of EGFP expression for NR1 cells 7-days post-transfection following 4-days of puromycin selection for transgene enrichment using a pGLP2-based gene-trap. **(g)** Diagram of plasmid recombination assay. **(h)** Flow cytometric quantification of plasmid recombination assay when used to test Bxb1 fused a C-myc, Nucleoplasmin (NLP), or Bipartite SV40+C-myc (BP) NLS while driven by EF-1α, hPGK, CAGG, and CBh promoters. mNeonGreen MFI was calculated for mCherry+ (transfected) population. **(i)** Optimization of FAST-STEM recombination by flow cytometric quantification of EGFP transgene integration into NR1 FAST-STEM hiPSCs by Bxb1 construct engineering and **(j)** transfection ratios and expansion medias, x-axis represents ng of p53DD-expression vector included in transfection mixture (of the 500 ng total DNA mixture transfected into a 24-well plate). **(k)** Representative fluorescence microscopy of EGFP expression for NR1 cells 5-days post-transfection with the optimized FAST-STEM delivery protocol **(l)** Tri-lineage differentiation marker and **(m)** transgene expression flow-cytometric quantification. **(n)** Diagram of dual-gene integration of EGFP and tagBFP transgenes selected for using puromycin and blasticidin respectively. **(o)** Flow cytometric quantification of EGFP and tagBFP transgene delivery into NR2 cells using dual puromycin/blasticidin selection, in addition to excision of the parental NR2 mScarlet cassette. **(p)** distribution of log_2_(skew) between barcode enrichment in the N_20_ plasmid library compared to enrichment of barcodes isolated from genomic DNA. skews lying outside |log_2_skew|≤2 are plotted in green, while those lying within nominal range are plotted in blue. **(q)** gini index of barcodes from the initial plasmid library compared to those isolated from transgene-delivered NR1 cells.

### Functional Screening of Regular FAST-STEM Integration in hiPSCs

To determine whether FAST-STEM cells could be effectively targeted for transgene delivery, flow cytometric quantification of an inserted fluorescent protein cassette was conducted. Due to the double-recombination event required for full integration of the FAST-STEM within the *AAVS1* locus, coupled with the homozygosity of the NR2 and NG2 cells, we theorized that many potential states could exist following gene delivery (Supplementary Figure 2a). Both dropout of the original FAST-STEM fluorescent marker to the delivered transgene, as well as the double-positive population were quantified for NG2 and NR2 FAST-STEM cells (Supplementary Figure 2b). Following ddPCR analysis of the double-positive and transgene-only populations for both NG2 and NR2, it was determined that both double-positive samples exhibited Neomycin copy-numbers greater than one, while transgene-only samples resulted in copy-numbers less than one but greater than 0. indicating this initial test only achieved incomplete removal of the FAST-STEM construct following gene delivery (Supplementary Figure 2c). Time from transfection to differentiation day 0 was 21 days – comparable with current landing-pad approaches.

### Co-transfection of Cytoprotective Factors During FAST-STEM Transgene Delivery Rescues Cell Viability and Transgene Expression

A significant increase in the degree of cell-death, reduction in live-cell population and reduced transfection efficiency was observed 48 hrs after co-transfection of an tagBFP transgene with pEF1α-Bxb1-NLS plasmids in NR1 cells compared to the transgene-only control (Figure 2b,c). Such an effect was even observed prior to removal of ROCK inhibitor from culture 1-day post-transfection (not-shown). We reasoned that transient double-strand break formation at the FAST-STEM recombination sites mediated by Bxb1 may have induced a cytotoxic response in hiPSCs. As such, co-transfection of a pGLP2-EF1a-tagBFP transgene, Bxb1 expression vector and plasmids encoding p53DD (a dominant negative fragment of the *p53* protein), Bcl-xL or XIAP were utilized to mute this response and rescue transgene expression^62,63^ (Figure 2b,c).

### ‘Fast-mode’ Transgene Delivery for Accelerated Design-Build-Test-Learn in hiPSCs

While initial proof-of-concept of ‘standard-mode’ recombination was functional, low efficiency and incomplete transgene removal inspired a more advanced ‘fast-mode’ mechanism for gene integration. Based on the promising result that cell viability during FAST-STEM transgene delivery could be rescued by co-transfection with cytoprotective factors, the efficiency of transgene delivery was quantified following 10-days post-transfection of NR1 cells with an EGFP transgene with no selective pressure (Figure 2d). Because no EGFP signal was detected in the no-Bxb1 control cells, we surmised any persisting EGFP fluorescence observed in test-samples was due to transgene delivery to the *AAVS1* locus. A marked increased transgene delivery efficiency (~336X) was observed in NR1 FAST-STEM hiPSCs when co-transfected with p53DD as opposed to an empty vector control, with a ~562X improvement compared to a simulation of the STRAIGHT-IN platform. A parallel experiment involving a 4-day puromycin selection regimen initiated 3-days post-transfection further demonstrates the rapidity of this system versus the current standard. The recombination-conditional puromycin-resistance conferred by this system allowed for an end-to-end 7-day transgenesis, making design-build-test-learn cycle time mush shorter than any other method (Figure 2e). Unlike initial attempts, a fractional amount of retained mScarlet expression (0.12% of total population, ~5.4% of EGFP+ population) was detected in cells transfected with p53DD supplementation, indicating near-complete excision of the original FAST-STEM reporter construct (Figure 2d). Furthermore, to ensure that supplementation with p53DD did not result in substantial spurious integration of transgene into the hiPSC genome, we tested whether inclusion of p53DD and/or Bxb1 during transfection with an EGFP transgene resulted in stable expression in both parental iPS11 and NR1 genetic backgrounds (Figure 2f). It was determined that on-target integration in the NR1 line was ~40X more efficient than off-target integration in the iPS11 parental line, indicating a 97.5% sample purity following transgenesis.

### Optimization of FAST-STEM Transgene Delivery

To further improve the transgenesis efficiency of the FAST-STEM hiPSC system, the effects of fusing Bxb1 to various nuclear localization signal (NLS) sequences and promoters were tested. The set of NLSs tested included C-myc, Nucleoplasmin (NLP), and a bipartite SV40+C-myc (BPNLS) derived from the PE2 prime editor while set of promoters included EF-1α, hPGK, CAGG, and CBh^64^. Initially, a plasmid-based recombination assay was implemented (Figure 2g). This system relied on recombination of an inverted mNeonGreen coding sequence to into the forward-orientation by a Bxb1-encoding plasmid (pBxO) such that recombination resulted in green fluorescence. The pBxO10 (CBh–Bxb1–C-myc) construct afforded a 2.2X improvement over the initial pBxO2 construct utilized herein (EF1a–Bxb1–NLP) (Figure 2h), and was highly promoter-dependent. Yet, such an effect was not replicable when an EGFP transgene was delivered into NR1 cells using this Bxb1 expression vector, resulting in a substantial reduction in transgenesis rate (Figure 2i). When all Bxb1 expression constructs were tested in this manner, it was determined that pBxO8 (CAGG–Bxb1–NLP) exhibited the highest recombination rate (383X higher than original transfection formulation) and minimized incomplete-RMCE resulting in retained mScarlet expression (Supplementary Figure 2d). This effect, however, was not significant when compared to some other Bxb1 construct variants. Furthermore, it was then determined that by combination of p53DD-based transient p53-inhibition, an optimal Bxb1 expression-construct, tuning the Bxb1:transgene:p53DD ratio, and media supplementation with eTeSR, a total improvement of 612X was achievable over initial standards (Figure 2j). Furthermore, retainment of the FAST-STEM reporter vector, indicating incomplete recombination, was minimized to ~2.7% of transgene-positive cells, indicating near-complete RMCE (Supplementary Figure 2e). This optimized protocol for FAST-STEM transgenesis was capable of achieving a selected monolayer within 5-days post-transfection (Figure 2k), a requirement for initiation of many differentiation protocols.

### Transgene Delivery to FAST-STEM hiPSCs does not Compromise Differentiation Potential

Finally, we demonstrated that NR2 cells engineered with an EGFP transgene which were FACS-sorted for EGFP+/mScarlet+ (termed NrG cells) using both standard (no CPG supplementation) and fast (optimized p53DD-supplemented formulation) transgenesis modes were able to differentiate efficiently into all three germ layers when compared to the iPS11 control. Furthermore, we found that these cells maintained transgene expression albeit with some context-dependent silencing (Figure 4l,m).

### Development of engineering modes that enhance design-build-test-learn capability

Given the exponentially longer timescales involved in applying design-build-test-learn engineering principles to stem cells, it would be valuable to obtain as much information per cycle as possible – i.e. optimize the “learn” step. To facilitate this, key innovations in cell engineering were adapted to this system, allowing engineers to: (i) deliver two different gene circuits simultaneously (ii) construct gene circuit libraries;

To accomplish dual gene circuit integration, and thus controlled copy-number of integrated transgenes, in homozygous NR2 cells, dual-selection with both puromycin and blasticidin was conducted to enrich for cells exhibiting both EGFP and tagBFP expression (Figure 2n,o). As well, we aimed to prove this system as a robust library-screening tool. A plasmid library of random barcodes (N_20_) was integrated into the genome of NR1 cells, and all barcodes with an initial abundance of ≥50 reads were tracked in sequence-data extracted from FAST-STEM cells. No detectable dropout of barcodes from the library after delivery was detected, with >95% of barcodes exhibiting difference in the ratio between the plasmid library barcode abundance and that of barcodes isolated from genomic DNA (skew) remaining within 2 log_2_-folds of one-another (Figure 2p). Furthermore, both the plasmid library and that isolated from FAST-STEM cells exhibited Gini indices ≤ 0.1 (0.034 – plasmid library, 0.089 – genomic library), a heuristic indication of sgRNA library uniformity (Figure 2q)^65^.

### Differentiation of FAST-STEM hiPSCs into clinically relevant terminal lineages

We next aimed to confirm that the FAST-STEM hiPSC lines were able to differentiate into specific cell lineages with the same robustness as the iPS11 parental line. Both the NR2 FAST-STEM hiPSC as well as an EGFP transgene-transfected NR2 sample FACS-sorted for both EGFP+/mScarlet+ phenotype (NrG) were directed towards six distinct cell lineages including hepatocytes (Figure 3a), neurons (Figure 3b), glial cells (Figure 3c), pancreatic beta-cells (Figure 3d), cardiomyocytes (Figure 3e) and skeletal muscle (Figure 3f).

**Figure 3:**
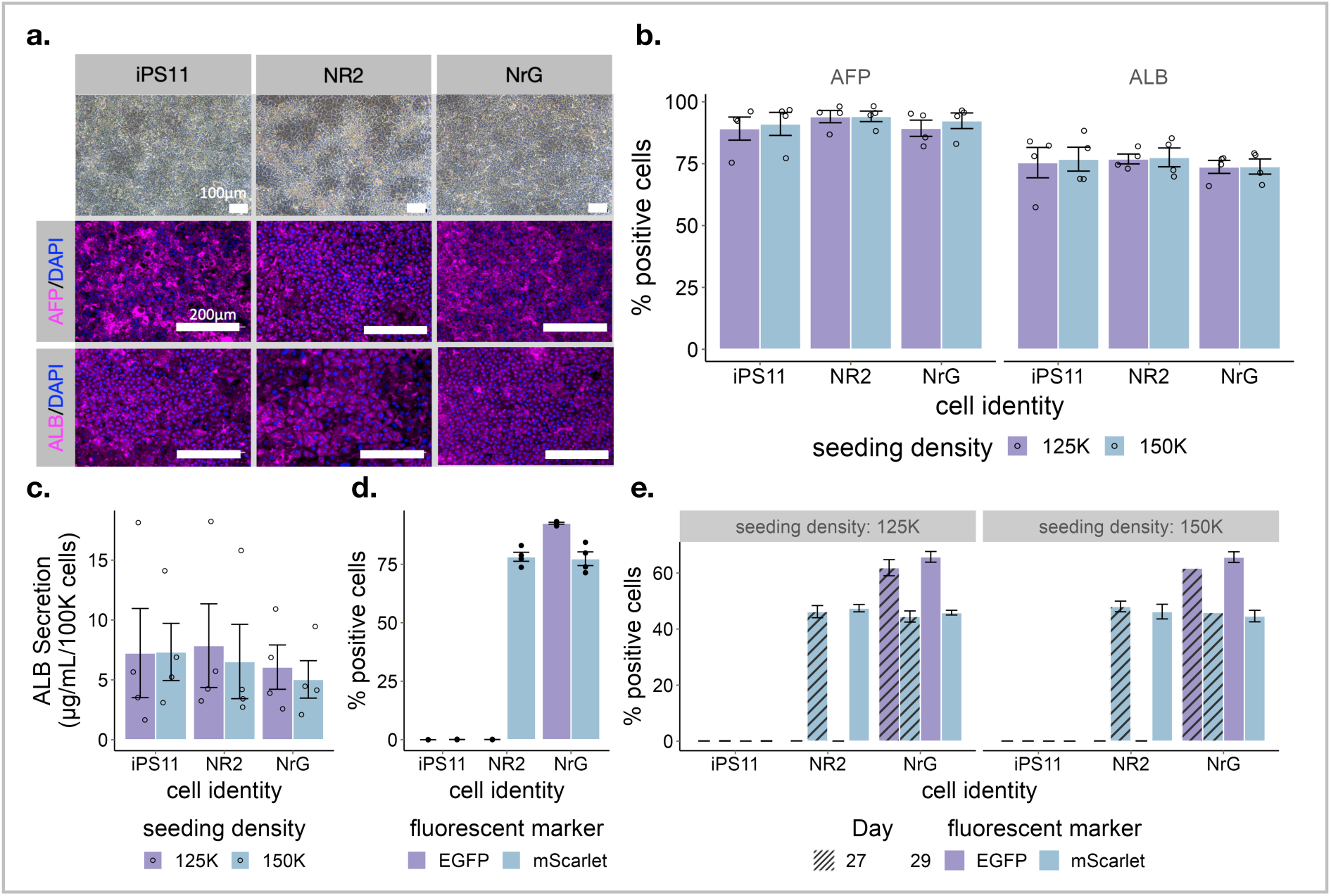
Differentiation of FAST-STEM hiPSCs into hepatoblasts. **(a)** Representative phase contrast images of hepatoblasts differentiated from iPS11, NR2, and NrG cell lines at day 29 (Scale bar, 100μm). In addition to representative immunofluorescence staining showing hepatoblasts expressing ALB and AFP at the end of differentiation (Scale bar, 200μm). **(b)** Quantification of the percentage of ALB+ and AFP+ hepatoblast populations at day 29 of differentiation with different seeding densities. **(c)** The levels of albumin secreted by hepatoblasts at the end of differentiation. The secretion level was detected by ELISA assay. **(d)** Quantification of GFP+ and mScarlet+ populations of IPS11, NR5 and NRGC1 at pluripotent stem cell and **(e)** differentiated hepatoblast stages.

#### Hepatoblast Differentiation

To assess the efficiency of hepatic progenitor cell differentiation (hepatoblast; HBs), iPS11, NR2, NrG, induced pluripotent stem cell lines were differentiated into HBs using a previously published protocol^66^. The commitment of definitive endoderm lineage was assessed through flow cytometry analysis of CXCR4 and cKIT at both day 7 and day 9 endoderm stages, indicating successful endoderm induction in all three cell lines (Supplementary Figure 3a). Remarkably, all three cell lines exhibited the generation of over 70% ALB+ and 90% AFP + cells from day 9 endoderm cells following 20 days for hepatic specification and differentiation (Figure 3a, Supplementary Figure 3b-d). Intriguingly, we do not see a significant impact of the initial cell density on the efficiency of HBs. The differentiated cells derived from all three iPS cell lines consistently displayed the typical cuboidal morphology characteristic of hepatic epithelial cells, which stained positive for ALB and AFP in immunostaining analyses (Figure 3a,b). Furthermore, we evaluated the hepatic function, particularly the secretion of human albumin into the culture medium, using ELISA assays. HB cells from all iPS cell lines exhibited comparable levels of human albumin secretion, quantified at 7.3±4.8 µg/mL for iPS11 cells, 7.9±7 μg/mL for NR2, and 6.1±3.7 μg/mL for NrG cells over a 48-hour duration, respectively (Figure 3c). During differentiation from the pluripotent stem cell stage to the hepatoblast stage, the population of mScarlet+ cells within NR2 exhibited a discernible decrease, declining from 78% to 46%. Similarly, NrG displayed a comparable pattern, with mScarlet+ populations diminishing from 77% to 44%. Notably, in the case of NrG, the population of EGFP+ cells also experienced a reduction, decreasing from 92% to 61% (Figure 3d,e).

#### Neural Precursor and Cortical Neuron Differentiation

To investigate the neural differentiation efficiency of the parental iPS11, NR2 and transfected-NR2 ‘NrG’ lines, the respective hiPSCs were provided with differentiation cues to generate neural progenitor cells (NPCs). Separately, we generated induced neurons (iNs) through over-expression of neurogenin-2, a transcription factor promoting rapid terminal neuronal differentiation (Figure 4a). NGN2 was delivered polyclonally in a lentiviral vector. The differentiation was assessed by RT-qPCR and immunostaining for known lineage specific markers levels such as Pax6 and Nestin for NPCs and MAP2 for iNs (Figure 4b-d). Importantly, the NR2 and NrG cell lines differentiated like their parental iPS11 line (Figure 4b,d). Furthermore, we monitored the mScarlet and EGFP transgene expression in the differentiated cells. 79.4 % of the starting NR2 hiPSCs cultures were mScarlet^+^ and we discerned no significant change in this proportion in the resulting NPCs and iNs (Figure 4d,e). The starting NrG hiPSC culture was composed of 49.8 % EGFP^+^/mScarlet+, 30.8 % EGFP^+^/mScarlet^−^, 18.4% EGFP^−^/mScarlet^−^, and 1.1% of EGFP^−^/mScarlet^+^ population (Figure 4d,e). There was no change in relative proportions of transgenic subpopulations’ composition during NPC generation. However, the proportions were significantly altered in induced neurons. Indeed, the resulting culture of iNs contained 10.5% of EGFP^−^/mScarlet^−^, 77.6% EGFP^+^/mScarlet^+^ cells, cells and 11.8% EGFP^+^/mScarlet^−^ cells (Figure 4d,e).

**Figure 4:**
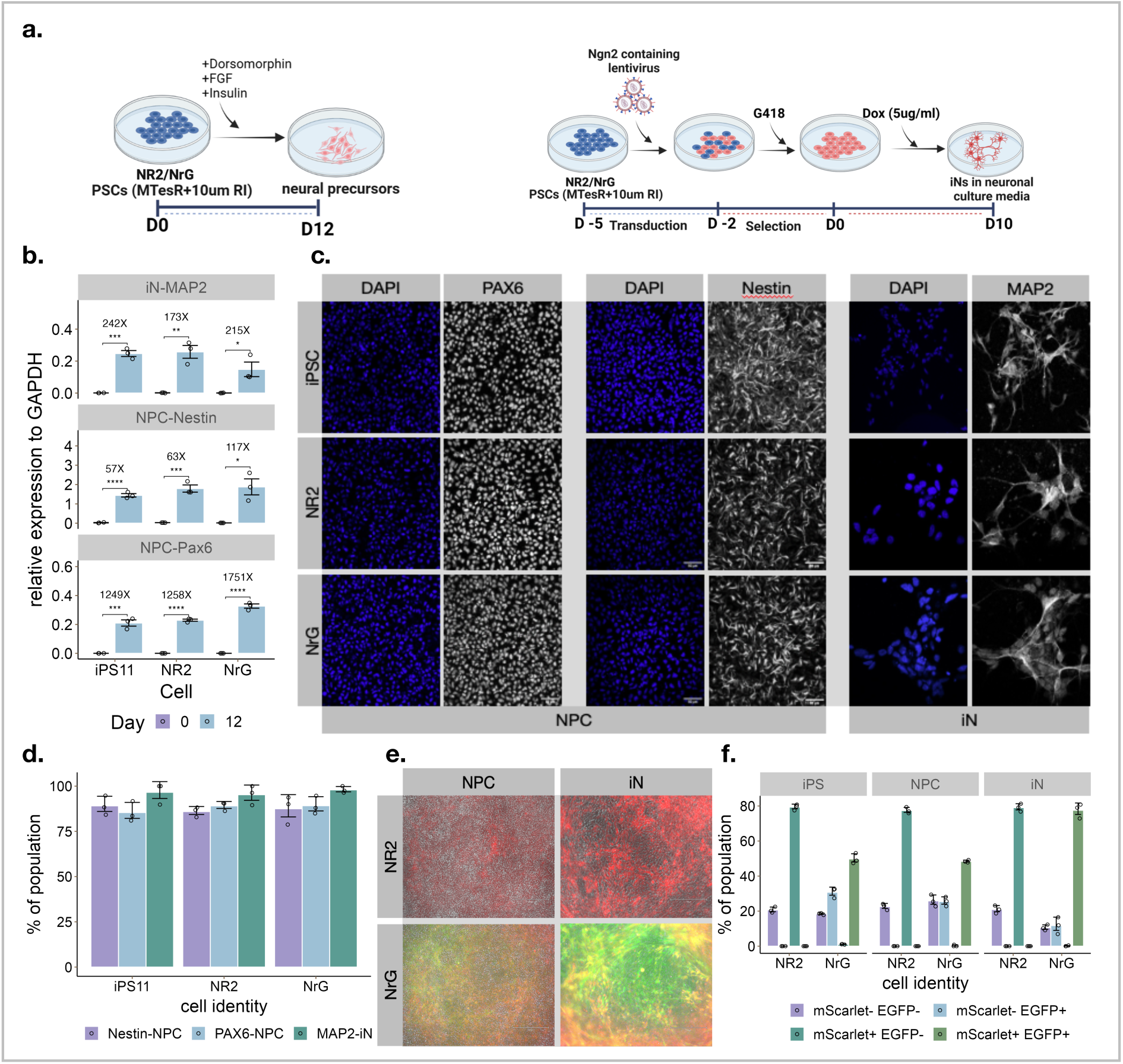
Differentiation of FAST-STEM hiPSCs into neural progenitors and cortical neurons. **(a)** Schematic of differentiation of hiPSCs to NPCs and iNs **(b)** RT-qPCR quantification of PAX6 and Nestin expression in NPCs and MAP2 in iNs for iPS11, NR2 and NrG cells at day 0 and day 12 of differentiation. **(c)** Representative immunostaining of PAX6 and Nestin during NPC-differentiation and MAP2 during iN-differentiation respectively for iPS11, NR2, and NrG cells and **(d)** quantitative image analysis thereof **(e)** Representative fluorescence microscopy of NR2 and NrG hiPSCs differentiated to NPC and iN lineages respectively. **(f)** transgene (EGFP) and FAST-STEM reporter (mScarlet) expression for hiPSC, NPC and iN lineages.

#### Glial Cell Differentiation

The parental iPS11, NR2 and transfected-NR2 ‘NrG’ lines were subjected to microglia-like cells (MLC) differentiations by providing hiPSCs with hematopoietic differentiation cues (Figure 5a). We confirmed the cell identity after MLC differentiation of iPS11, NR2 and NrG hiPSC by RT-qPCR using known lineage specific markers such as AIF1, CD11B, CX3CR1, HEXB, and TGFBR1 (Figure 5b). The relative gene expression of these markers was significantly higher in differentiated MLC lineage as compared to day 0 in all three cell lines indicating that all NR2 and NrG cell lines differentiated in as similar manner as that of parental iPS11(Figure 3c-iii). We investigated the efficient differentiation of NR2 and NrG cell lines to MLC by counting the percentage of cells positive for IBA1 and PU.1 to those of populations produced by the parental iPS11. (Figure 5c,d). In differentiated iPS11, NR2 and NrG MLCs, >97% population were IBA1^+^ and >80% were PU.1^+^. Furthermore, we manually quantified the proportions of cells positive for mScarlet and/or EGFP transgene expression on epifluorescent images through the differentiation process. The starting NR2 and NrG hiPSC cultures were the same as described above (Figure 4f). Following differentiation, NR2 MLCs were 18.9% mScarlet^+^. Meanwhile, NrG MLCs were 33.8% EGFP^−^/mScarlet^−^, 37.4% EGFP^+^/mScarlet^−^, 23.5% EGFP^+^/mScarlet^+^ and 5,3% EGFP^−^ mScarlet^+^ (Figure 5e)

**Figure 5:**
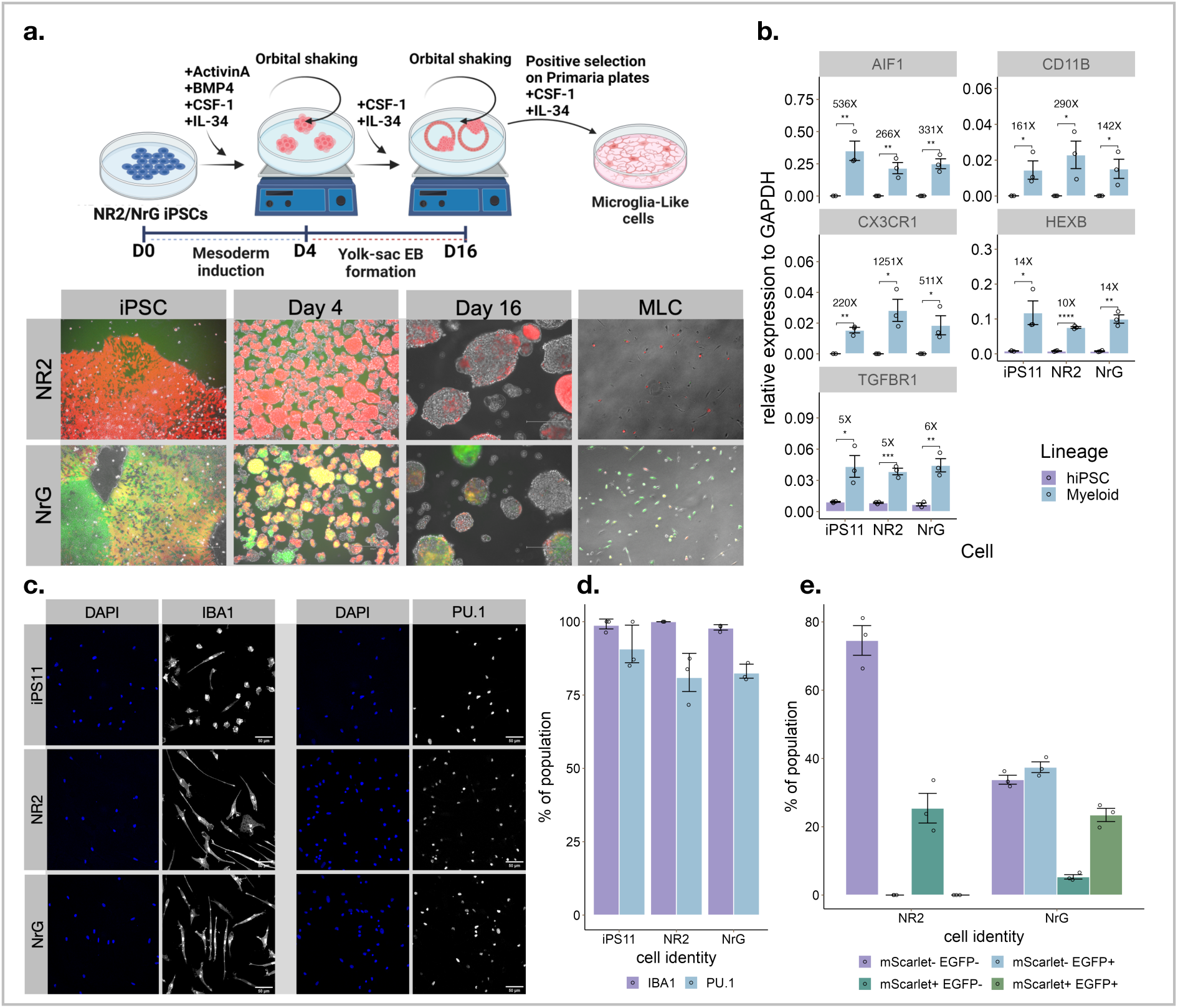
Differentiation of FAST-STEM hiPSCs into microglia-like cells (MLCs). **(a)** Differentiation schematic and representative fluorescence microscopy of NR2 and NrG hiPSCs through discrete phases of differentiation. **(b)** RT-qPCR quantification of AIF1, CD11B, CX3CR1, HEXB, and TGFBR1 for MLC-differentiation of iPS11, NR2, and NrG cells. **(c)** Representative immunostaining of IBA1 and PU.1 respectively for iPS11, NR2, and NrG cells differentiated to MLCs and **(d)** quantitative image analysis thereof. **(e)** transgene (EGFP) and FAST-STEM reporter (mScarlet) expression for hiPSC and MLC lineages.

#### Islet-like Cell Differentiation

iPS11 and its derivative cell lines (NR2, and NrG) were differentiated to islet-like cells using published protocols (Fig. 6a)^67–69^. No significant differences were observed in the differentiation of definitive endoderm as shown by the frequency of CXCR4+/CKIT+ and SOX17+/FOXA2+ cells obtained at day 3 of differentiation (Supplementary Figure 3d – i-iv). Each lines gave rise to similar frequencies of pancreatic progenitor cells (PDX1+/NKX6-1+) at day 12 (Figure 6b), early beta-like cells (C-PEP+/NKX6-1+) at day 23 (Supplementary Figure 3e), and late beta-like cells (C-PEP+/NKX6-1+) at day 29 (Figure 6b). mScarlet+ and EGFP+ expression was monitored throughout differentiation by flow cytometry analysis (Figure 6c) and (Supplementary Figure 3f). While mScarlet expression remained consistent throughout differentiation, the frequency of EGFP+ cells were significantly lower from day 8 onwards compared to day 0 undifferentiated cells.

**Figure 6:**
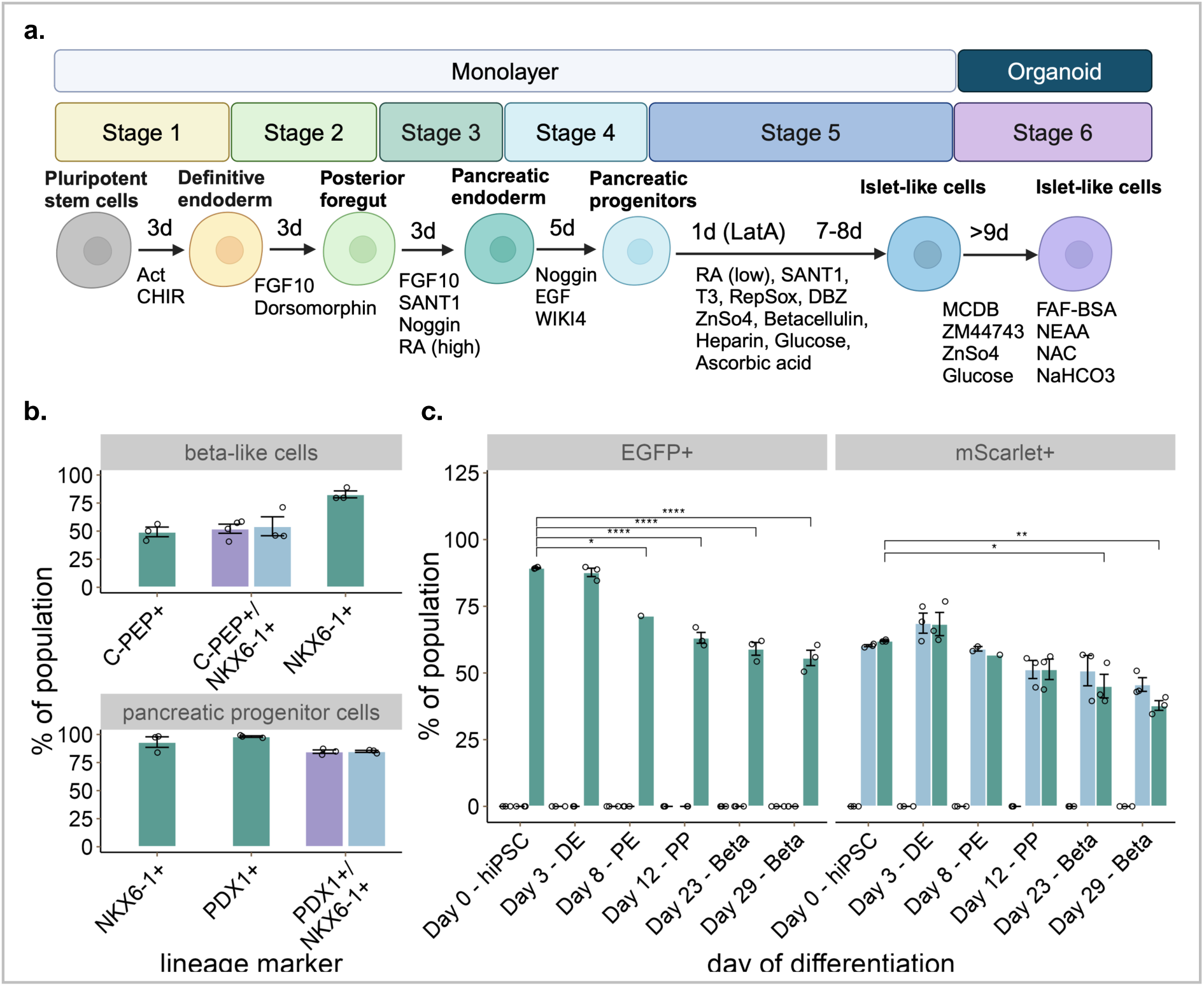
Differentiation of FAST-STEM hiPSCs into islet-like cells. **(a)** Schematic representation of pluripotent stem cells differentiation into islet-like cells, **(b)** Pancreatic progenitor marker PDX1+/NKX6-1+ and Beta-like cells marker C-PEP+/NKX6-1+ expression analysis by flow cytometry (n=3 independent cell differentiations) at the end of stage 4 (day 12). All data are represented as the mean, and all error bars represent the standard error of mean. **(c)** mScarlet and EGFP reporter expression during multiple stages of stem cell differentiation to islet-like cells. Time points of differentiation are shown as day 0 (hPSC), day 3 (definitive endoderm - DE), day 12 (pancreatic progenitors - PP), day 23 (Islet-like cells - Beta) and day 29 (Islet-like cells - Beta).

#### Cardiomyocyte Differentiation

We optimized a previously established 20-day cardiomyocyte differentiation protocol for the parental iPS11, NR2 and NrG lines (see Methods) ^70,71^. All three lines successfully generated populations of >50% PDGFRα+ CD56+ mesoderm cells at day 3 (Fig 7a,b), which gave rise to populations of >70% cTnT+ cardiomyocytes at day 20 (Fig 7c,d). We measured expression of the EGFP and mScarlet transgenes in the NrG line during cardiomyocyte differentiation (Fig 7e,f) and observed moderate silencing of both transgenes (Fig 7g). At the end of differentiation, the total proportion of EGFP+ cells had declined from 87% ± 1% at day 0 to 70% ± 2% at day 20 (~0.2-fold) while the total proportion of mScarlet+ cells had declined from 59% ± 1% at day 0 to 51% ± 1% (~0.14-fold). Consequently, there was also a ~0.24-fold decrease in the proportion of cells co-expressing EGFP and mScarlet (55% ± 1% at day 0 to 41% ± 1% at day 20).

**Figure 7:**
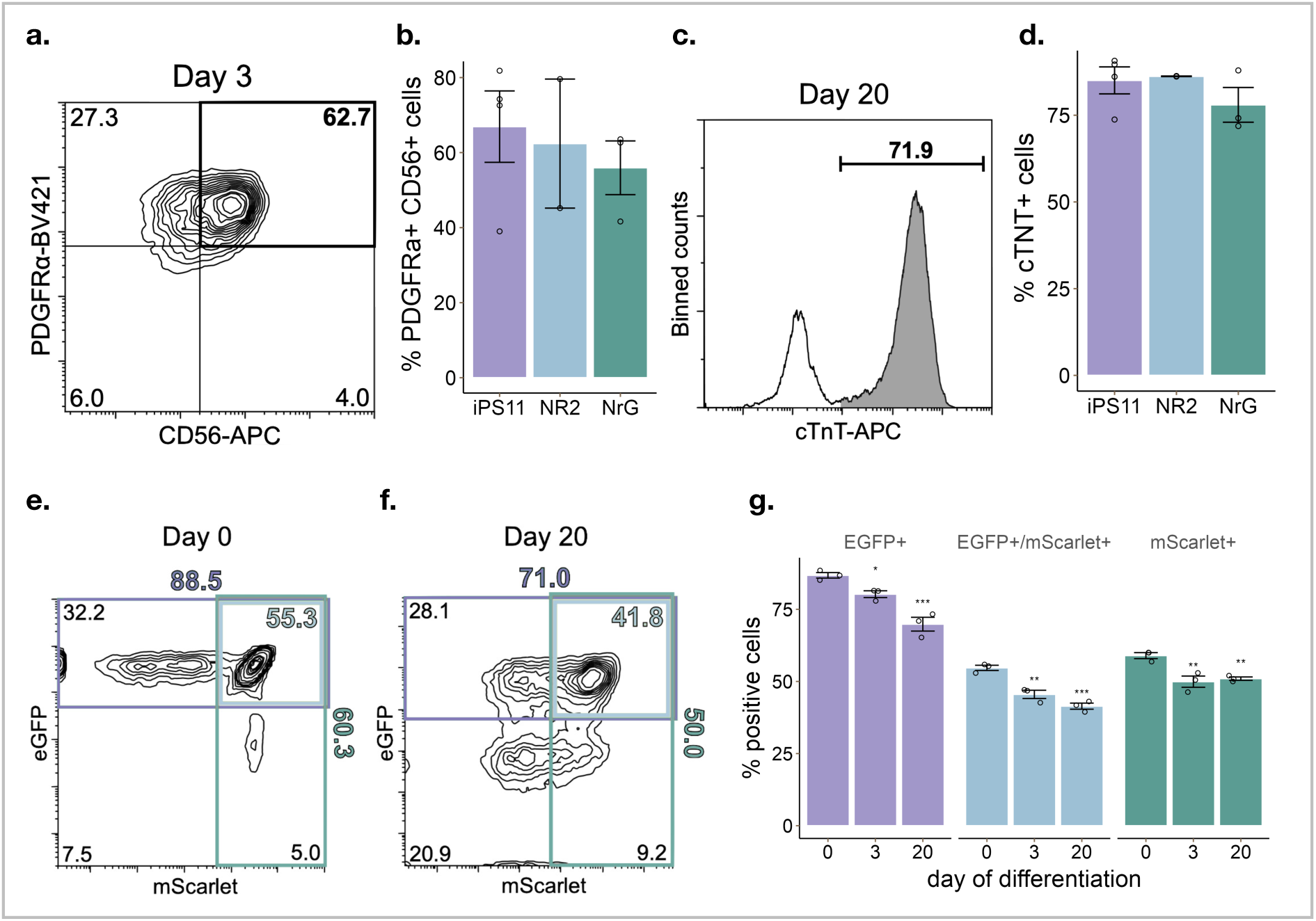
Differentiation of FAST-STEM hiPSCs into cardiomyocytes. **(a)** Analysis of FAST-STEM transgene expression during cardiomyocyte differentiation. Representative flow cytometric analysis of the proportion of cells expressing mesoderm markers PDGFRα and CD56 at day 3 of differentiation. **(b)** Average proportion of PDGFRα+ CD56+ mesoderm cells for iPS11, NR2 and NrG at day 3 of differentiation (one-way ANOVA not significant p = 0.744, iPS11 n = 4, NR2 n = 2, NrG n = 3). **(c)** Representative flow cytometric analysis of the proportion of cells expressing the cardiomyocyte marker cTnT day 20 of differentiation. **(d)** Average proportion of cTnT+ cardiomyocytes for iPS11, NR2 and NrG at day 20 of differentiation (one-way ANOVA not significant p = 0.416, iPS11 n = 4, NR2 n = 2, NrG n = 3). **(e, f)** Representative flow cytometric analyses of eGFP and mScarlet transgene expression in NrG cells at **(e)** day 0 and **(f)** day 20 of differentiation. **(g)** Average proportion of total eGFP+ (green), total mScarlet+ (red) and double EGFP+ mScarlet+ (yellow) cells at day 0, day 3 and day 20 of differentiation (one-way ANOVA with Dunnett’s post-hoc test comparing to day 0 condition for each transgene group, n = 3). Error bars represent SEM. Statistical significance: *** p < 0.001, ** p < 0.01, * p < 0.05.

#### Skeletal Muscle Differentiation

The parental iPS11, NR2 and NrG cells were directed to the skeletal muscle myogenic progenitor lineage using a previously reported 35-day method^35^. Flow cytometric analysis of the passage 2 cell populations revealed a similar frequency of CD56+ myogenic progenitors (Figure 8b) with <1% HNK-1 contaminating cells (data not shown) across all of the cell lines. We observed that 74.8 ± 0.7% of passage 2 NR2-derived myogenic progenitors were mScarlet+, while the NrG-derived population was found to be only 39.2 ± 0.9% mScarlet+ and 70.8 ± 0.4% EGFP+ (Figure 8c). Importantly, passage 2 myogenic progenitors derived from all lines produced phenotypically similar cultures of multinucleated, striated skeletal muscle myotubes when placed in low serum conditions (Figure 8a).

**Figure 8:**
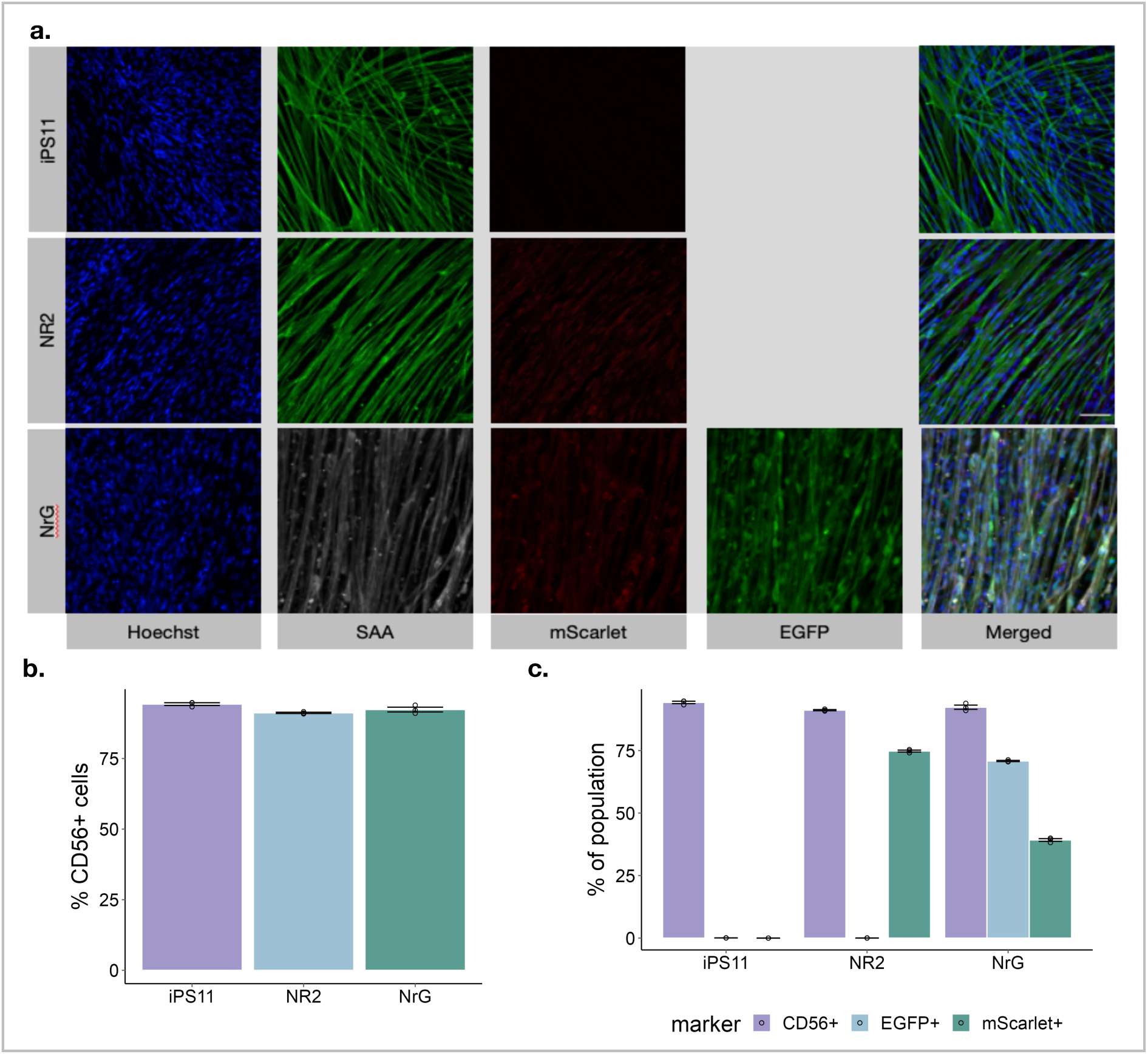
Differentiation of FAST-STEM hiPSCs into skeletal muscle. **(a)** Representative 20x confocal images of multinucleated skeletal muscle cells produced from iPS11, NR2, and NrG FAST-STEM hiPSCs immunostained to visualize nuclei (Hoechst), sarcomeric alpha-actinin (SAA), mScarlet, and EGFP. **(b)** CD56 staining of iPS11, NR2, and NrG FAST-STEM cells on day 35 of skeletal muscle differentiation. **(c)** Fluorescent marker expression of mScarlet (parental FAST-STEM reporter) and EGFP transgene in hiPSC-derived skeletal muscle.

## Discussion

Despite much fanfare, integrating synthetic biology with clinically relevant hiPSC-derived cell types has proven exceedingly challenging to execute. The platform presented here is our foundational contribution toward dismantling barriers that currently preclude cell-based therapy prototyping and development using this approach. We envisage this system in a manner akin to breadboard for electrical engineers, amenable to flexible and rapid changes without the laborious effort of synthesizing a microchip for every design. It builds on previous work and attempts to address previously identified limitations. While extant landing-pad systems have effectively delivered genes to hiPSCs, the use of Cre recombinase as the core gene-transfer effector may lead to reduced accuracy and efficiency when compared to Bxb1, and in some cases pluripotency and differentiation potential were not validated^72^. Others utilize the ϕC31 LSR as a core effector^73^. This enzyme exhibits substantial permissiveness for chromosomal rearrangements of the human genome and imprecise recombination in comparison to Bxb1^72,74,75^. As such, the use of RMCE catalyzed by Bxb1 alone may provide a safer, less off-target prone, transgene delivery to FAST-STEM hiPSCs. Furthermore, the use of a splice-acceptor gene trap in *AAVS1* driven by the endogenous *PPP1R12C* promoter can facilitate antibiotic resistance conditional on FAST-STEM targeting, enabling rapid isolation of recombined hiPSCs^76^.

The approach presented here uses a Bxb1-based RMCE-compatible construct placed into the *AAVS1* safe-harbour locus using CRISPR-HDR in hiPSCs. While both plasmid and long-single-stranded-DNA (lssDNA)-based HDR donors were utilized to knock-in cells, this study presents the first lssDNA-based donor strategy in hiPSCs using chemical transfection to our knowledge, which confirms previous reports of this method outperforming plasmid-based CRISPR-HDR in other systems^77^. It is anticipated that this method will be an effective tool for stable transfection with reduced costs compared to use of a nucleofector, simplifying and democratizing cell-line generation and potentially reducing associated cytotoxicity during transfection.

An in-depth slate of experiments for verification of FAST-STEM hiPSC validity was conducted (Figure 1) to ensure that the cells developed herein would be capable of downstream use. PCR of the *AAVS1* locus containing the FAST-STEM construct, followed by nanopore-sequencing leveraged a recent trend in cost-effective whole-plasmid and PCR product sequencing. Investigators can use similar methodologies to confirm transgene delivery with sequence-level certainty. Confirmation of pluripotency of the NB1, NR1, NG2, and NR2 hiPSC clones (Figure 1b), as well as differentiation into all three germ layers (Figure 1c-e) validated them for general use. Transgene expression was maintained to varying degrees in these lineages (Supplementary Figure 1e) and was highly dependent on fluorescent protein, with tagBFP exhibiting drastic silencing in comparison to EGFP and mScarlet. Determination of the normal karyotype (Supplementary Figure 1f-i) of both lines was the confirmed that genetic manipulation of these cells did not adversely affect their genomic integrity. Integration of transgenes did not impact downstream differentiation potential to all three germ layers in either normal or fast recombination-modes (Figure 2l,m).

During initial functional screening of NG2 and NR2 cells to confirm transgene delivery (Supplementary Figure 2b), it was observed that mixed populations of seemingly homozygous or heterozygous could be isolated, expressing either both transgene and original FAST-STEM fluorescent reporter, or only transgene (Supplementary Figure 2b). Yet, after ddPCR analysis of reporter (Neomycin resistance gene) copy number (Supplementary Figure 2c), it was hypothesized that the phenotypes observed were potentially caused by transgene silencing or incomplete removal of the FAST-STEM construct. As such, it was determined that improving the efficiency of the recombination reaction in hiPSCs was highly desirable. Reducing apoptosis due to genomic manipulation-induced cytotoxicity has been an effective method for improving the efficiency of CRISPR HDR and prime editing by expression of p53DD and upregulation of Bcl-xL in hiPSCs without inducing long-term genomic instability^62,78,79^. As such, it was theorized that transiently decoupling the DNA-damage and/or apoptosis signaling axes following transfection could have a similar effect. It was confirmed that hiPSCs experience a substantial reduction in viability and transgenesis following transfection with Bxb1, which could be ameliorated by supplementation with p53DD (Figure 2b-e) without substantial introduction of off-target integration events (Figure 2f). This approach also substantially reduced the rate of incomplete removal of the FAST-STEM construct following RMCE (Figure 2d, Supplementary Figure 2d,e) to a minimum (~2.5%).

Bxb1 constructs were further optimized by promoter and NLS engineering, finding that while the plasmid-based recombination assay (Figure 2f,g) identified a CBh-driven Bxb1 expression vector fused to a C-myc NLS as most efficient, such results were not replicable when transfecting NR1 cells (Figure 2h). Optimization with all Bxb1 constructs in NR1 cells showed that CAG-driven Bxb1 with a nucleoplasmin NLS was most efficient at precipitating recombination in this system. This suggests that during the initial recombination assay, where numerous pRA3 plasmids could be localized in cytoplasm, recombination was not representative of a single-target configuration. The latter is more akin to the FAST-STEM system where nuclear localization of both delivery vector and Bxb1 are obligatory for recombination. This suggests that nuclear recombinase expression is the rate-limiting factor for ideal stable transgene integration.

To contextualize the need for the >600-fold improvement in recombination rate over current standards for hiPSC engineering demonstrated herein, a typical synthetic biology project can require between tens to hundreds of design-build-test-learn cycles. In the age of generative-AI-augmented biomedical discovery, this number may trend towards the later figure. Because gene knock-in, selection, monoclonal expansion, screening, banking, and testing of engineered hiPSCs takes months and substantial financial investment in order to achieve a uniform population, we foresee simplifying this process to a working-week (5-days) as a paradigm-shifting development. Current methods are not scalable to parallelized engineering of numerous gene circuits or candidate sequences into hiPSCs, where the workload for developing more cells is roughly linear with respect to time. For example, engineering an entire 96-well plate of distinct gene circuits into hiPSCs accurately whilst retaining differentiation potential would be logistically impossible by current standards within a reasonable timeframe, yet could be achievable with relative ease using FAST-STEM. Utilizing this innovation therefore has the potential to reduce typical stem cell engineering project timelines by between six months to many years. In many instances it may prove the difference between a project being feasible or not.

Broad utility of FAST-STEM as a regenerative medicine/stem cell engineering tool, was demonstrated by an ecosystem of nine stem-cell laboratories, that were able to successfully direct differentiation of transgene-loaded hiPSCs into six disparate lineages with similar efficiency to the iPS11-control sample (Figures 3-8). Silencing of the FAST-STEM mScarlet reporter and to a lesser extent, the EGFP transgene, was observed at variable magnitudes across lineages. Epigenetic silencing in stem cells is an ongoing issue in the field and work to solve it will be enabled and catalysed by this tool. Further investigation is required to alleviate this in a lineage-specific manner. Suggested directions may include promoter choice and engineering, CpG-depletion, or incorporation of insulators, matrix-attachment regions and upstream-chromatin-opening elements into expression vectors.

Delivery of multiple transgenes is an invaluable tool for synthetic biologists to develop increasingly complex and intelligent cell therapies. Applications include, but are not limited to, rapid prototyping, combinatorial gene expression, and isolation of genetic circuits to reduce cross-talk. While such a capability has been demonstrated in a limited context in transformed mammalian cells, this is the first demonstration of simultaneous and controllable integration into two parallel loci efficiently with full semi-scarless excision of delivery plasmid backbone without the need for numerous fluorescent reporters, kill-switches, and FACS-sorting (Figure 2n,o)^80,81^. FAST-STEM is capable of simultaneous selection with puromycin/blasticidin to strictly enforce copy-number, which is invaluable for delivery of two distinct gene circuits, or for testing the effect of transgene copy-number on cell function if one transgene is replaced with an empty vector. The ability to conduct such an experiment sequentially further bolsters the FAST-STEM system for use as an iterative design-build-test-learn tool, as an initial experiment using hiPSCs expressing a given transgene may direct the choice for subsequent integration of a second gene circuit.

Library screening is classically accomplished using lentivirus as a means of infecting cells with single-copies of gene variants or sgRNAs of interest^65,82^. This method, while effective for transducing cells at-scale has numerous key flaws^83^. Firstly, due to the diploid-ssRNA genome of lentiviridae, it is theoretically impossible to truly infect cells at the single-copy level. Furthermore, it is highly likely that two non-identical library elements are present on each packaged ssRNA molecule respectively, further complicating the readout of lentiviral screens. Secondly, while researchers generally transduce libraries into cells at a lentiviral multiplicity-of-infection (MOI) of 0.3X in order to achieve a nominal single gene copy in most cells, such an effect is stochastic, with a small (~10%) proportion of cells achieving >1X MOI. Thirdly, recombination of lentiviral elements by template-switching frequently results in barcode mismatches with sequences of interest^83–85^. Finally, random-integration of the lentivirus into the host genome may introduce heterogeneity and inconsistency in screening results, as positional effects may modulate library element expression. As such, landing-pad systems such as FAST-STEM present an alternative route for library-delivery which alleviates these concerns by enforcing a defined copy-number in all cells^86,87^. at a validated safe locus. A 1581-element degenerate barcode (N_20_) plasmid library was delivered to NR1 cells which maintained its diversity following integration (Figure 2p,q). Utility of the FAST-STEM system in this manner may dramatically improve confidence and reproducibility in library-screening experiments conducted in hiPSCs or differentiated lineages thereof.

In addition to the key applications of FAST-STEM described herein. Two classically-laborious procedures in hiPSC engineering may be facilitated by this platform. For example, generating stable gene-specific shRNA knock-down lines can be particularly challenging, yet the highly repeatable expression of transgenes from the FAST-STEM system may vastly accelerate the time to generate such cells, enabling facile disease modeling. Modulation of gene expression using the many components of the CRISPR toolkit could be an effective use-case for the FAST-STEM system. Canonically, such experiments may be cumbersome due to the large payload size and heterogeneous expression of Cas9-based effectors in hiPSCs, but may be overcame by this system.

FAST-STEM is a multifaceted and accessible tool, intended to facilitate––and dramatically accelerate––projects that involve the engineering of human stem cells and derived tissues. The primary goal is for use in next-generation cell-based therapy development, where integration of synthetic biology and generative A.I. has the potential for development of cures for a high number of diseases. Success of the design-build-test-learn approach (the ‘secret sauce’ for many other branches of synthetic biology) has not yet been effectively leveraged by stem cell engineers, due to the extremely long timelines that stem cell work involves. FAST-STEM is a big step toward reducing those timelines toward enabling rapid development of human cell based therapeutic devices. Beyond cell-based therapy, FAST-STEM could also be deployed in contexts including disease modelling and biomaterials development. It provides facilities for sophisticated experimental strategies involving successive transgene integrations, copy number control, library screens, and any combination thereof.

## Methods

### General Plasmid Construction and Molecular Cloning

Polymerase chain reaction (PCR) of constructs utilized for plasmid assembly were amplified by NEB *Q5* Hot-Start 2X Mastermix according to the manufacturer’s protocol (New England Biolabs, Ipswitch, MA, USA; M0494L). Amplifications of difficult templates (such as the CAG promoter) were conducted using Platinum SuperFi II Green PCR Master Mix (Invitrogen, Waltham, MA, USA; 12369010). PCR reactions were purified using the Monarch DNA PCR Cleanup Kit (New England Biolabs; T1030). For Gibson assembly of constructs, *NEBuilder* HiFi 2X Mastermix (New England Biolabs; E2621L) was utilized according to the manufacturer’s protocol^88^. All assembly reactions were transformed into homebrew NEB *Stable E. coli* (New England Biolabs; C3040) made using the Mix&Go transformation kit (Zymo Research, Irvine, CA, USA; T3001) and grown at 30°C to ensure construct stability. Plasmid clones were picked and inoculated into liquid LB media for overnight culture followed by miniprep using the PureLink Plasmid MiniPrep Kit (Invitrogen; K210010), and verified using sanger-sequencing by Eurofins genomics (Louisville, KY, USA). All plasmids generated in this study are documented in Supplementary Table 1.

### FAST-STEM Donor and Cas9/gRNA Expression Construct Assembly

In order to construct each donor vector, the backbone of pSH-EFIRES-P-AtAFB2-mCherry (Addgene, Watertown, MA, USA; 129716) was PCR-amplified to include both homology arms to *AAVS1*. Both Bxb1 *attP*(GA) and *attB*(GT) sites were synthesized by oligonucleotide overlap assembly as designed by *Primerize*^89,90^. Other fragments encoding the EF1α promoter, EGFP, mScarlet, P2A, and neomycin resistance were PCR-amplified from extant constructs present in our laboratory using *Q5* Hot-Start 2X Mastermix. HDR-donor assemblies were completed using NEBuilder HiFi 2X Mastermix and verified by whole-plasmid sequencing by Primordium Labs (Arcadia, CA, USA).

An all-in-one Cas9/gRNA expression construct encoding an *AAVS1* sgRNA sequence derived from pCas9-sgAAVS1-2 PX458 (Addgene; 129727) was placed downstream of the hU6 promoter in a PX458 (Addgene; 48138) Cas9 expression vector by golden-gate assembly with BbsI-HF (New England Biolabs, R3539)^91,92^. Following bacterial transformation, clonal expansion, and plasmid screening by sanger sequencing, correct plasmid clones were further expanded and isolated using the ZymoPure II Plasmid MaxiPrep Kit (Zymo Research; D4203).

### Long Single-stranded DNA (lssDNA) HDR-Donor Generation

The lssDNA generation protocol was adapted from Rosenstein and Walker^93^. Briefly, PCR amplification of each donor construct and flanking *AAVS1* homology arms was conducted using a forward primer modified with a 5’-biotin moiety and five 5’-phosphorothioate linkages, and a 5’-phosphorylated reverse primer. 200 μL reactions using Herculase II DNA polymerase (Agilent, Santa Clara, CA, USA; 600675) were conducted according to the manufacturer’s protocols. PCR reactions were purified and digested with 10U of lambda exonuclease (New England Biolabs; M0262) in 1X lambda-exonuclease buffer for 2 hours at 37°C, followed by addition of 10 U dsDNAse (Thermofisher Scientific, Waltham, MA, USA; EN0771) for 10 min, and heat inactivation at 65°C for 20 min. Reactions were purified according to the ssDNA cleanup protocol from the Monarch PCR cleanup kit.

### hiPSC Culture

iPS11 hiPSCs (Alstem Cell Advancements, Richmond, CA, USA; iPS11) cultured with mTeSR-Plus (StemCell Technologies, Vancouver, BC, Canada; 100-0275) according to the manufacturer’s protocol on GelTrex-coated plates (Invitrogen; A1413201). Unless otherwise stated, ROCK Inhibitor Y-27632 was added at 1X concentration for all seeding and passaging steps. Cells were grown in a 5% CO_2_ incubator set to 37°C. Passaging was conducted using TrypLE gentle-dissociation reagent according to standard tissue culture protocols (Invitrogen; 12604013). Cell cryopreservation was conducted by resuspending hiPSCs in CryoStor CS10 (StemCell Technologies; 100-1061) freezing media followed by fixed-rate cooling using a Mr. Frosty Freezing Container (Thermo Scientific; 5100-0001).

### CRISPR-HDR Mediated Knock-In of FAST-STEM Donor DNA in hiPSCs

hiPSCs were grown to 60% confluence in a 24-well plate, and media was changed and supplemented with 1X RevitaCell 30 min prior to transfection with a 500 ng of a 1:1 mass-ratio of PX458:lssDNA donor for each FAST-STEM construct using lipofectamine stem in accordance with the manufacturers protocol (Invitrogen; STEM00001).

### Selection and Clonal Expansion of FAST-STEM hiPSCs

3-days post-transfection, cells from each well were dissociated and replated at 30% confluency, followed by supplementation of culture media with 50 μg/mL G418 (Invitrogen; 10131035), which was increased by 50 μg/mL daily until reaching a final concentration of 250 μg/mL for 10 days of routine passaging, facilitating enrichment of the population of hiPSCs stably-transfected with the FAST-STEM vector. Cells were then dissociated and resuspended in cloning media consisting of mTeSR-Plus supplemented with 1X CloneR2 (StemCell Technologies; 100-0691) and mixed gently with a P1000 pipette followed by passing through a 40 μm cell strainer in order to generate a single-cell suspension. Cell samples were counted using a Countess II Haemocytometer (Life Technologies, Carlsbad, CA, USA), and plated onto single wells of a 6-well tissue culture plate at a density of 50 cells/cm^2^ with 2 mL of cloning media. 2 days later, culture media was changed with fresh cloning media. On the 4^th^ day, cloning media was removed and replaced with mTeSR-Plus supplemented with 250 μg/mL G418 selective antibiotic. Once discrete 200-300 cell colonies were visible, eight colonies per-well were picked, dissociated, expanded, and split into duplicate 24-well plates.

### Verification of FAST-STEM Construct Insertion

While one of the 24-well plates containing all FAST-STEM hiPSC clones was maintained for expansion, FACS-sorting for the fluorescent reporter, and cryopreservation, the second was used for gDNA extraction using the Invitrogen PureLink Genomic DNA Mini Kit (Invitrogen; K182001). PCR was conducted using primers flanking the *AAVS1* homology arms (AAVS1_screenF: ACCTGCCCAGTACAGGCATC, AAVS1_screenR: TCCCCGTT GCCAGTCTCGAT) present on the FAST-STEM donor construct. PCR was conducted with Phire II Hot-Start PCR 2X mastermix (Thermo Scientific; F125L) using 100 ng of gDNA template, 250 nM of AAVS1_screenF/R primers respectively and 3% v/v DMSO and was run at an initial denaturation temperature of 98°C for 30 sec, followed by 35 cycles of 98°C for 15 sec, 68°C for 20 sec, and 72°C for 2:45 min, and finished by a 2 min final extension at 72°C. Following PCR cleanup, agarose gel electrophoresis was conducted to confirm correct integration into the *AAVS1* site and an initial readout of zygosity. ddPCR was conducted by the Center for Applied Genomics (TCAG, Toronto, ON, Canada) in order to calculate the copy-number of FAST-STEM constructs in each candidate cell line by absolute quantification of the neomycin resistance gene normalized to the human RNase P gene.

### Fluorescence-Activated Cell Sorting (FACS) of hiPSCs

hiPSC clones which passed all verification steps were grown to 80% confluency in 6-well plates and were dissociated with 750 μL of TrypLE. Following incubation at 37°C for 5 mins, 750 μL of mTeSR Plus was added, and cells were pelleted by centrifugation at 200xg for 5 mins. Cells were resuspended in sorting buffer (mTeSR Plus supplemented with 1X CloneR2) at a density of 4 million cells/mL, and were pipette-mixed vigorously to dissociate cell aggregates. Cells were filtered through a 40 μm nylon cell-strainer into a 5 mL polystyrene FACS tube. A collection tube of 2 ml mTeSR-plus supplemented with 1X CloneR2 was prepared for deposition of sorted cells. Cells were sorted on a FACS Aria IIIu by staff members of the University of Toronto - Temerty Faculty of Medicine Flow Cytometry Facility staff members. NB cells were gated on tagBFP+ expression (Pacific Blue Filter, 405 nm laser), NG cells were gated on +EGFP expression (FITC filter, 488 nm laser), while NR cells were gated on +mScarlet expression (PE-Cy5 filter, 561 nm laser). Cells were cultured for 4-days undisturbed post-sorting before downstream manipulation and/or cell-banking.

### Flow Cytometric Quantification of hiPSC Pluripotency Markers

Staining for *OCT-4, SOX-2, and NANOG* was conducted with the following primary antibodies respectively: anti-Oct-3/4 (BD Biosciences, San Jose, CA, USA), rabbit anti-Sox-2 (New England Biolabs), and rabbit anti-NANOG (New England Biolabs). Briefly, cells were detatched with TrypLE (Invitrogen) and quenched with mTeSR Plus, followed by a 5 min centrifugation at 200xG for 5 minutes and by a wash-step with PBS. Cells were then incubated with respective primary antibodies at room temperature for 1 hour, followed by two PBS-wash steps and secondary antibody staining and analysis on an LSRFortessa flow cytometer (BD Biosciences), which was also used for all downstream flow cytometry.

### Differentiation of hiPSC to Endoderm, Mesoderm and Ectoderm and Confirmation Thereof

Endoderm differentiation was adapted from the stage-1 protocol described in Nostro *et. al.,* with cells maintained in d1-media until day-4 of differentiation^94^. Cells were then dethatched, PBS-washed, and split for subsequent endodermal marker staining. A third of each cell sample was stained for 30 mins at 4°C with 1:100-dilutions of APC-conjugated anti-*C-Kit*/CD117 (BD Biosciences; 550412) and PE-Cy7-conjugated anti-*CXCR-4*/CD184 (BD Biosciences; 560669), followed by two washes with PBS+5% FBS.

Mesoderm differentiation was adapted from the cardiomyocyte differentiation protocol of Li et al. and Dubois et al. and assessed on day-4^71,95^. A day prior to induction, an Aggrewell-400 plate (StemCell Technologies; 34411) was coated with 1 mL of 5% Pluoronic F-68 (Gibco, Waltham, MA, USA; 24040032) and incubated for 30 minutes at room temperature before replacement with PBS for storage. On the first day of differentiation, FAST-STEM hiPSCs grown to 80% confluency were dissociated into single cells using TypLE for 5 minutes and seeded into the Aggrewell at a density of 1.2×10^5^ cells per-well in 1 mL of mTesR Plus with 10 μg/mL ROCK Inhibitor Y-27632 (Tocris Bioscience, Bristol, UK; 1254) and 1% Penicillin-Streptomycin (Life Technologies; 15140122). The following day (day 2), media was replaced with 800 µl of StemPro-34 SFM (Gibco; 10639011) supplemented with 0.5% Transferrin (Roche, Basel, Switzerland; 10652202001), 0.1% Ascorbic Acid (Millipore Sigma; A8960), 0.3% MTG (Millipore Sigma; M6145), 10 ng/µL BMP4 (R&D Systems, 314BP), 5 ng/µL bFGF (PeproTech, 100-18B) and 6 ng/mL Activin A (StemCell Technologies; 78001) for commencement of stage-1 induction. On day 4, the media was removed carefully and aggregates were washed with non-supplemented StemPro before dissociation with TrypLE for flow-cytometry preparation. In brief, cells were costained with a 1:50 dilution of anti-CD140a (BD Biosciences; 556001) primary antibody and a 1:50 dilution of Brilliant-Violet-conjugated anti-CD56 (BD Biosciences; 562780) in 50 µL of PBS + 5% FBS for 30 minutes followed by a 1:1000 dilution of Alexa Fluor 488-conjugated secondary antibody (Invitrogen; A10631) in 50 µL of PBS + 5% FBS for another 30 minutes.

Ectoderm differentiation was conducted using the STEMdiff™ SMADi Neural Induction Kit (StemCell Technologies; 08581) according to the manufacturer’s protocol. On day-8 and/or 16 of differentiation, cells were detatched, PBS-washed, and fixed with 4% PFA for 15 mins on ice, followed by permeabilization with ice-cold absolute methanol for 3 mins. Cells were then washed with PBS+5% FBS and stained with 1:100-dilutions of AF647-conjugated anti-PAX-6 (BD Biosciences; 562249) and PerCP-Cy5.5-conjugated anti-SOX-2 (BD Biosciences; 561506).

### Assembly of FAST-STEM Delivery Vectors and associated transgenes

Plasmid DNA assembly was completed using NEBuilder HiFi 2X Mastermix. *Primerize* was used to design *attB* and *attP* assemblies^89,90^. a pTwist-Amp-High-Copy backbone was utilized and assembled with ALOXE3 and cHS4 insulators PCR amplified from hiPSC gDNA and MTK1_009 (Addgene #123672) respectively to form the pGLP0,1 and 2 constructs. Delivery vectors were verified by whole-plasmid sequencing by Primordium Labs.

Cloning of transcriptional units (TUs) was conducted in accordance with MTK toolkit procedures using NEBridge Golden Gate BsaI Assembly Mastermix (New England Biolabs; E1601) according to the manufacturer’s protocol^61^. Part identifiers used for assembling all TU vectors are itemized in Supplementary Table 2. TUs were assembled into pGLP-class destination vectors using NEBridge Golden Gate BsmBI Assembly Mastermix (New England Biolabs; E1602) according to the manufacturer’s protocol, and itemized in Supplementary Table 3.

### Initial Functional Screening of FAST-STEM Integration Efficiency

In order to quantify FAST-STEM hiPSC recombination efficiency, the *NG2* and *NR2* cell lines were transiently co-transfected with pEf1a-Bxb1-NLS and pGLP2-based delivery vector (pGLP2-Ef1a-mCherry for NG, pGLP2-Ef1a-EGFP for NR) with Lipofectamine stem in a 3:1 plasmid mass ratio. As well, control reactions excluding the Bxb1-integrase expression vector were ran to determine when transient transfection of delivery vector had subsided. ~48 hrs post-transfection, 0.25 ug/mL puromycin selective antibiotic (Millipore Sigma, Burlington, MA, USA; P7255) was added to culture, and media was changed daily until no live-cells were visible in the no-integrase control well. Cells were dissociated with TrypLE, and replated for further expansion and downstream FACS. ddPCR of transgene-loaded NG2 and NR2 cells with conducted by TCAG.

### Improving Transgene Delivery Potential of FAST-STEM hiPSCs

For detection of the effect of p53DD, Bcl-xL, and XIAP on transfection and recombination, 2×10^5^ NR1 hiPSCs per-well were plated in a 12-well plate in triplicate using standard culturing procedures described above. The following day, cells were transfected with either transgene only (400 ng pGLP2-EF1α-tagBFP + 600 ng pUC19), transgene with Bxb1 (400 ng pGLP2-EF1α-tagBFP, 400 ng pEF1a-Bxb1-NLS, 200 ng pUC19), or transgene with Bxb1 and one of the three cytoprotective genes (400 ng pGLP2-EF1α-tagBFP, 400 ng pEF1a-Bxb1-NLS, 200 ng pEF1a-[p53DD/Bcl-xL/XIAP]). The following day, media was changed to mTeSR without ROCK inhibitor supplementation (standard practice 1-day post-transfection). 48-hours post-transfection, cells were dissociated and stained with LIVE/DEAD fixable near-IR dead-cell stain (L10119, Invitrogen) and immunostained with PerCP/Cy5.5-conjugated anti-Annexin V (Biolegend, San Diego, CA, USA; 640935). Finally, cells were analyzed by flow cytometry with appropriate compensation controls for viability (LIVE/DEAD), transgene expression (tagBFP), apoptotic initiation (annexin-V), and FAST-STEM reporter expression (mScarlet).

For initial gene-delivery testing, plating and transfections were conducted in an identical manner described above with the following modifications: an EGFP-transgene was used instead of tagBFP and cultures were scaled-down based on surface area by a factor of two plating in 24-well as opposed to 12-well plates. In addition, a simulation of the STRAIGHT-IN platform was added (200 ng pGLP2-PuroR-EGFP, 300 ng pCAG-NLS-HA-Bxb1) was added. 48-hours post-transfection, cells were dissociated with accutase (StemCell Technologies; 07920) and replated into a 96-well plate at a 1:5 dilution (accounting for 1:5 reduction in surface area between 24 to 96-well plate, thus conserving cell density) and preserved without selection until transgene expression in the no-Bxb1 control was no longer visible, followed by cell dissociation and quantification of transgene expression and excision of the FAST-STEM reporter.

For fast gene integration at 5-days. Cells were plated at 100K cells/well in a 24-well plate and supplemented with eTeSR and ROCK inhibitor, followed by transfection as described above. One day post-transfection, media was changed to fresh eTeSR and supplemented with 0.5 μg/mL puromycin. On the 3^rd^ day post-transfection, cells were lifted and replated into a 96-well plate, followed by a media change the day after. Cells were then imaged at using an Incucyte SX5 live-cell imager (Sartorius, Göttingen, Germany) to track confluency and EGFP transgene expression.

Off-target integration assessment of this system was conducted by transfection of either iPS11 or NR1 FAST-STEM hiPSCs with 175 ng of pGLP2-PuroR-EGFP with or without 175 ng of pEF1a-Bxb1-NLS vector and p53DD expression vector respectively. Cells were propagated without antibiotic selective pressure for 14-days post-transfection and analyzed by flow cytometry for EGFP expression.

While the initial pEF1α-Bxb1-NLS construct utilized an EF-1α promoter and Nucleoplasmin NLS at the C-terminus, numerous Bxb1 expression constructs were assembled using golden gate cloning as depicted in Supplementary Table 3. The pRA3 plasmid was constructed using Gibson assembly to place an mNeonGreen fluorescent protein between Bxb1 attP and attB into an hPGK-promoter based backbone. All pBxO-class Bxb1 plasmids were co-transfected using lipofectamine stem into WT iPS11 cells plated at 40% confluency in 24-well plates and supplemented with ROCK Inhibitor Y-27632 such that each well received 200 ng pBxO, 100 ng p53DD, 100 ng pRA3 and 100ng pEF1a-mCherry transfection marker. Compensation controls were also transfected for mNeonGreen, mCherry. Cells were cultured for three days followed by dissociation, viability staining with LIVE/DEAD-Near IR, and flow cytometric analysis.

Quantification of Bxb1 expression construct efficiency by direct measurement using the FAST-STEM system was conducted in an identical manner to that described above (co-transfection of pGLP2-EGFP, p53DD, and Bxb1 vectors followed by expansion and flow cytometry). Finally, optimizing transfection conditions for transgene delivery was conducted using varying ratios of p53DD expression vector (1:5, 3:10, 2:5 of total DNA by mass) the optimized Bxb1 expression vector (pBxO8) at varying ratios with transgene (1:3, 1:1, 3:1 by remaining mass following supplementation with p53DD vector). Furthermore, ROCK inhibitor was replaced by CloneR2 to improve post-transfection viability and growth-rate. Where indicated, transfection and cell expansion was conducted using eTeSR (StemCell Technologies; 100-1215), an optimized medium for single-cell expansion of hiPSCs to further improve growth rate during cell engineering.

### Optimized Transfection Protocol for FAST-STEM hiPSC Gene Delivery

Because of the rapid post-transfection recovery and growth of FAST-STEM hiPSCs using optimized conditions for ‘fast mode’ gene delivery, we developed an optimized passaging regimen summarized as follows which prevents cells prematurely reaching confluency (thus reducing selection efficiency and growth rate). On day-1, Cells were seeded at a density 1×10^5^ cells/well in a 24-well plate using eTeSR supplemented with 1X CloneR2, and transfected the day after (day 0) with a DNA mixture of 100 ng p53DD vector, 100 ng pBxO8, and 300 ng pGLP2 delivery vector. On day 1, media was replaced for un-supplemented eTeSR, and cells were passaged with accutase into one well of a 6-well plate on day 2 with CloneR2 supplementation. On day 3, media was replaced with eTeSR+0.5 μg/mL puromycin. On day 5, cells were passaged back into a well of a 24-well plate with both CloneR2 and puromycin supplementation. On day 6, media was refreshed with puromycin supplementation. By day 7, a confluent monolayer of engineered hiPSCs should be achievable. While we found this protocol to work best in our hands, our anecdotal experience is that this protocol is exceedingly flexible, and cells can be cultured without increasing surface area on the day-2 passage by reducing initial seeding densities, or using mTeSR Plus instead of eTeSR to slow growth kinetics. This will, however, result in a lower cell yield by day-7.

### Dual gene delivery into NR2 FAST-STEM hiPSCs

Transfections were completed using the optimized transfection protocol described above. For dual-transgene loading with puromycin/blasticidin selection, pGLP2-PuroR-EF-1α-EGFP and pGLP2-BlastR-EF-1α-tagBFP were transfected in an equimolar ratio. Selection was conducted using 0.25 μg/mL puromycin and 1 μg/mL blasticidin S antibiotics.

### Library Screening in NR1 FAST-STEM hiPSCs

A random barcode library was assembled by primerize-based assembly of a gene fragment encoding a random (N_20_) segment with golden-gate-compatible overhangs for assembly into the pGLP2-PuroR delivery vector. A 50 μL BsmBI-based assembly reaction was conducted according to standard NEB protocols, and transformed into NEB Stable *E. coli.,* followed by a 90 minute outgrowth and plating onto 4 150mm LB agar plates. The following day, colonies were pooled by addition of 10 mL of cold LB broth per-plate, scraped with a bacterial plate spreader, and plasmid DNA was extracted using the ZymoPure II Plasmid MaxiPrep Kit. Transfection was conducted in three wells of a six-well plate. Cells were expanded and selected with puromycin (0.25 μg/mL) and gDNA was extracted with a PureLink Genomic DNA Mini Kit. PCR using primers specific for the barcoded region of the FAST-STEM delivery vector with Genewiz Amplicon-EZ-compatible overhangs was conducted for both the initial plasmid library and isolated gDNA library, with secondary barcodes denoting library identity. Samples were submitted for 2×250 Illumina paired-end amplicon sequencing using Genewiz Amplicon-EZ, ~700,000 reads were generated.

### Library screen analysis pipeline

Barcode content was extracted and quantified from sequencing reads. Barcodes with abundances ≥50 reads in the original plasmid library were further tracked in the genomic DNA barcode library to ensure faithful quantitation of library performance. Enrichment of each barcode in both plasmid and genomic libraries were determined, and skew was calculated as the ratio of plasmid/genomic enrichment scores. Gini indices were calculated for both libraries using log-transformed counts of barcode reads, as indicated in Li et. al.^65^

### Differentiation of hiPSC to Hepatoblasts

hiPSCs were cultured with StemMACS™ iPS-Brew XF (Miltenyi; 130-104-368) according to the manufacturer’s protocol on GelTrex-coated plates. To start hepatoblast differentiation, the hiPSCs were dissociated with TrypLE and passaged to 12-well plates coated with Matrigel (Corning, Corning, NY, USA; 354234) with a seeding density of 125000/well and 150000/well. Hepatoblast differentiation was adapted from the protocol described in Ogawa et al^66^.

### Flow Cytometric Analysis of Hepatoblast Differentiation

To measure the fluorescent levels, hiPSCs and hiPSC-derived hepatoblasts were dissociated with TrypLE. GFP was detected by the FITC channel, while mScarlet was detected by the PE channel.

Day 7 or day 9 endoderm cells were dissociated into single cells with TrypLE for 5 minutes at 37°C. For surface marker analyses, cells were stained for 30 minutes at room temperature in FACS buffer consisting of PBS (Corning; 21-031-CM) with 5% FCS. The following antibodies were used: 1:50 dilution of anti-CXCR4-APC (BD Biosciences; BD555976), and 1:100-dilution of anti-cKIT-APC (Invitrogen; CD11705).

For intracellular staining, cells were fixed for 30 minutes at room temperature with 4% paraformaldehyde (Electron Microscopy Science, Hatfield, PA, USA; 15700) in PBS and then permeabilized with 90% methanol for 10 minutes at –30°C. The permeabilized cells were washed with FACS buffer and stained with unconjugated primary antibodies in FACS buffer for 1 hour at room temperature. The following antibodies were used: 1:200 dilution of goat anti-human Albumin (Bethyl Laboratories, Montgomerey, TX, USA; A80-129A), 1:2000 dilution of rabbit anti-human AFP (Agilent; A000829), isotype rabbit IgG (Jackson Immunoresearch, Westgrove, PA, USA; 001-000-003), and isotype goat IgG (Millipore Sigma; I5256). Cells were subsequently stained with secondary antibodies for 30 minutes at room temperature, a 1:400-dilution of both donkey anti-rabbit IgG-Alexa Fluor 647 (Invitrogen; A31573) and donkey anti-goat IgG-Alexa Fluor 647 (Invitrogen; A21447) were used. The stained cells were analyzed using an LSR Fortessa flow cytometer (BD).

### Immunostaining of Hepatoblasts

Hepatoblasts were fixed with 4% PFA for 20 mins at room temperature and permeabilized with cold 100% methanol for 10 mins. Cells were washed twice with wash buffer consisting of PBS and 0.1% BSA and blocked with DAKO blocking reagent (Dako: X0909) for 5 mins. Cells were stained with rabbit anti-human AFP (1:2000) and goat anti-human ALB (1:200) respectively in staining buffer consisting of PBS with 0.1% Triton X and 0.1% BSA overnight at 4℃. The stained cells were washed three times with wash buffer and then stained with donkey anti-rabbit IgG-Alexa Fluor 647 (1:400) and donkey anti-goat IgG-Alexa Fluor 647 (1:400) respectively for 1 hour at room temperature followed by DAPI (BIOTIUM: 40043) staining. The stained cells were analyzed using an EVOS microscope (Thermo Fisher).

### Hepatoblast Albumin Secretion Assay

The supernatants were harvested at the end of differentiation and the amount of albumin secreted was measured using the human albumin ELISA kit (Bethyl laboratories) according to the manufacturer’s protocol.

### Neural Precursor (NP) Cell Differentiation

iPS11, NR2 and NrG stem cells were differentiated to three independent neural precursor (NP) cells by culturing them with dorsomorphin, fibroblast growth factor (FGF) and insulin as previously described^86^. NPs were maintained in the NGD medium (Neuro-glial differentiation medium 0.5X, as previously described^85)^ containing 10ng/ml bFGF and 10ng/mL human insulin on matrigel-coated plates and passaged using accutase every 5-7 days.

### Neuronal Induction

iPS11, NR-5 and NrG stem cells were transduced separately three times by lentivirus containing empty backbone vector and neurogenin (ngn2) plasmid (pLV-TetO-ngn2-neo)^96^. The transduced cells were induced with 5µg/ml doxycycline for five days.

### Microglia-like cell (MLC) differentiation

iPS11, NR2 and NrG cell lines were differentiated into MLCs using a protocol modified from previous publication^84^. hiPSC colonies were treated with ReLeSR (STEMCELL, 05873) according to manufacturer’s protocol and triturated to form a suspension of uniform clumps with an average diameter of 100um, transferred directly to the NGD, supplemented with 50ng/mL hIL-34, 50ng/mL hCSF1, 50ng/ml hBMP4, 1ng/ml hActivin-A and 10uM Rock inhibitor Y26732 in ultra-low attachment 6-well plates (corning). One confluent hiPSC well was split into 6 suspension wells to achieve 30-40k spheroids/well density. After 24 hours, spheroids were collected and fed with fresh medium without ROCK inhibitor. BMP4 and Activin-A were removed after 4 days, and then NGD + IL-34 + CSF1 was replaced every 4 days. Embryoid bodies were monitored for exclusion of compact phase-bright neuralized spheroids, and expansion of cystic yolk sac embryoid bodies. From day 14 to 28 post differentiation, EBs were gently triturated to shear off loose cells of interest (primitive macrophages) every other day, settled to remove the EBs in a 15mL falcon tube, and the supernatant was placed in a single well of a Primaria 6-well plate. Cells were left to adhere overnight. Unattached cells and small EBs were washed with HBSS+0.1% BSA, and wells were fed with fresh NGD + IL-34 + CSF1. Attached cells were monitored for morphological characteristics of microglial precursors (compact nucleus, vacuoles, membrane ruffles, motility), and wells from 5-6 consecutive productions were pooled to constitute one mixed birthdate population. Three independent MLC differentiations were performed with iPS11, NR2 and NrG lines.

### RNA Extraction, Reverse Transcription and Quantitative PCR for Neuron and Microglial Cell Differentiation

Cells were homogenized and total RNA extracted using the RNeasy kit (Qiagen #74134) according to the manufacturer’s instructions. Total RNA concentrations were measured using NanoDrop ND-1000 spectrophotometer. RNA was reverse transcribed into cDNA using Superscript IV reverse transcriptase (Invitrogen, Thermo Fisher Scientific #18090200) with random hexamer primers according. Transcript abundance was determined by quantitative PCR using SYBR Green PCR mix (Applied Biosystems, Thermo Fisher Scientific #A25778). The primers were designed according to specific targets (Table 1). Raw Ct values were normalized to GAPDH.

### Immuno-Fluorescent Staining and Imaging of Neurons and Microglia

Cells were fixed with 4% (w/v) paraformaldehyde (Cedarlane Labs 15710(EM)) in PBS. Following membrane permeabilization with PBS containing 0.3% Triton x-100 (Sigma, T8787), cells were blocked with 3% normal donkey serum (Sigma, S30-M). Primary antibodies were against PAX6, NESTIN and MAP2 for neurons as well as PU.1 and IBA1 for microglia and visualized by secondary antibodies conjugated with Alexa 647 (Thermo) followed by counter-staining with DAPI (Thermo, D3571) (Table 2). Fluorescent images of immuno-staining were captured on a Nikon AR-1 confocal microscope.

### Cell Counting for Neuron and Microglia Differentiation

Cells were single cell dissociated and seeded sparse at 15k/cm^2^ (for MLC) 25k/cm^2^. Cells were allowed to settle prior to stain with 1ug/ml Hoescht for 15min. Cells were imaged using an EVOS M5000 imaging system (Invitrogen). Hoescht stained nuclei, mScarlet and EGFP labelled cells were counted using ImageJ.

### Differentiation into pancreatic islet-like cells *in vitro*

iPS11 and their transgenic derivatives cell lines such as NR2 and NrG were differentiated into islet-like cells by protocols based on published previously^67–69^ with some modifications as indicated below.

#### Day-1

iPS11 wild type (WT) and transgenic cell lines were grown in feeder-free conditions such as hiPSC Brew XF medium (Miltenyi Biotec) in Geltrex (Life technologies) coated 6 well plates (Falcon). Prior to start an experiment, cells were replated at density of ~200,000/cm^2^ to achieve confluency and start the differentiation process 24 hours after plating.

#### Stage-1: Definitive endoderm induction (DE) (Day 0 – Day 3)

The basal medium consists of MCDB131 (Wisent), Glutamine (1x) (Hyclone), NaHCO3 (1.5g/L) (Gibco), Fatty acid free BSA (0.5%) (Proliant), D-Glucose (5mM) (Sigma), Ascorbic acid (50μg/ml) (Sigma). *Day 0-*Cells were washed with 1X DPBS-/-followed by definitive endoderm induction medium were changed with Activin-A (100ng/ml) (R& D Systems), CHIR990210 (3μM) (Tocris Bioscience), ROCK Inhibitor Y-27632. *Day 1-*replace with definitive endoderm induction basal medium with Activin-A (100ng/ml) + CHIR990210 (0.3μM). *Day 2-*replace with definitive endoderm induction basal medium with Activin-A (100ng/ml) only.

#### Stage-2: Posterior foregut induction (PFG) (Day 3-Day 5)

The basal medium consists of MCDB131, Glutamine (1x), NaHCO3 (1.5g/L), Fatty acid free BSA (0.5%), D-Glucose (5mM), Ascorbic acid (50μg/ml) supplemented with FGF10 (50ng/ml) (R&D Systems) + Dorsomorphin (0.75μM) (Sigma). Medium was replaced on day 5.

#### Stage-3: Pancreatic endoderm induction (PE) (Day 6-Day 8)

The basal medium consists of DMEM-High Glucose (Gibco), Glutamine (1x), 1% vol/vol NeuroBrew-21 without Vitamin A (Miltenyi Biotec), Ascorbic acid (50μg/ml) supplemented with FGF10 (50ng/ml), hNoggin (50ng/ml) (R&D Systems), 0.25μM SANT-1 (Tocris bioscience) and 2μM all-trans retinoic acid (RA) (Sigma). Medium was replaced every day until day 8.

#### Stage-4: Pancreatic progenitors induction (PP) (Day 8-Day 12)

The basal medium consists of DMEM-High Glucose, Glutamine (1x), 1% vol/vol NeuroBrew-21 without Vitamin A, Ascorbic acid (50μg/ml) supplemented with hNoggin, hEGF(100ng/ml), (R&D Systems), WIKI4 (0.9μM) (Tocris bioscience) and medium was replaced every alternative day until day 12.

#### Stage-5: Endocrine cells differentiation (Beta-like cells) (Day 12-Day 23)

The basal medium consists of MCDB131, Glutamine (1x), NaHCO3 (1.5g/L), D-Glucose (15mM), 1% vol/vol NeuroBrew-21 without Vitamin A, 1μM T3 (3,3’,5-Triiiodo-L-thyronine) (Sigma), Ascorbic acid (50μg/ml) supplemented with Heparin (10μg/ml) (Sigma), 0.25μM SANT-1, 10μM RepSox (Tocris bioscience), 10μM ZnSo4 (Sigma), and 0.05μM all-trans retinoic acid (RA), 100nM DBZ (Tocris bioscience). 0.3μM Latrunculin A (LatA) (Tocris bioscience). After 24 hours, at day 13 medium was replaced without LatA and with BetaCellulin (R&D Systems). Subsequently same new media were replaced on day 15, day 17, day 20 and maintained until day 23.

#### Stage-6: Beta-like cells differentiation (Beta-like cells) (Day 23-Day 29+)

At day 23, cells were mechanically dissociated and cultured them in suspension cultures in regular Petri dishes. The basal medium consists of MCDB131, Glutamine (1x), NaHCO3 (1.5g/L), D-Glucose (1.5mM), 1% non-essential amino acids (NEAA)(Gibco), Fatty acid free BSA (0.5%), 10μM ZnSo4 supplemented with 1mM N-Acetyl Cysteine (NAC) (Sigma), 0.5μm ZM447439 (Selleckchem). The media were replaced every 2-3 days until day 29+.

### Flow Cytometric Qantification of Beta-Cell Differentiation

For intracellular staining, monolayer Cells and cell aggregates were dissociated into single cells using TryPLE Express Enzyme (Gibco) at 37°C for 3-5 minutes (monolayer) or 10-12 minutes (aggregates). The cells were washed and pelleted in PBS with 5% FBS and 10 μl/ml DNase (FACS buffer), filtered and stained with Zombie Violet (BioLegend) for Live/dead staining 1/1000 in PBS at room temperature for 20 minutes, washed and fixed the cells in BD-Cytofix/Cytoperm solution (BD Biosciences) at room temperature for 30 minutes and washed store them in 1X BD-PermWash (BD Biosciences) solution at 4°C until to begin the staining process. Flowcytometry analysis was done in multiple time points, Stage 1, definitive endoderm for live (CXCR4+/CKIT+)/ fixed (SOX17+/FOXA2+) at day 3, Stage 4, pancreatic progenitors (PDX1+/NKX6-1+) at day 12 and Stage 5, beta-like cells (C-PEP+/NKX6-1+) at day 23 and Stage 6, beta-like cells (C-PEP+/NKX6-1+) at day 29. The respective combination of primary antibodies was diluted in 1X Perm/Wash buffer and incubated overnight at 4°C. Next day, cells were washed and stained with secondary antibodies diluted in Perm/Wash buffer for 30 minutes at room temperature. Then, cells were washed and stored in FACS buffer for flow cytometry analyses. For cell surface marker staining (CXCR4/CKIT): At day 3 definitive endoderm live cells were incubated with directly conjugated antibodies in FACS buffer for 20 minutes at 4°C. After incubation, samples were washed twice and stained with DAPI before flow cytometry analyses. Stage specific markers and reporter expression (mScarlet+/eGFP+) were quantified using flow cytometry and data are analyzed using FlowJo software. (BD Biosciences). The list of primary and secondary antibody dilution details are listed in Supplementary table 1.1 and 1.2.

### Differentiation of hiPSCs to Cardiomyocytes

Cardiomyocyte differentiation was adapted from protocols established by Kattman et al. and Lee et al^70,71^. hiPSCs were cultured in mTeSR1 medium (StemCell Technologies) on hESC-qualified Matrigel and passaged using TrypLE according to standard tissue culture protocols. hiPSCs were cultured for at least 2 passages and grown to 80-90% confluency before differentiation. *Day 0 (aggregation):* hiPSCs were dissociated into single cells and aggregated to form embryoid bodies (EBs) in mTeSR1 medium supplemented with 10 μM ROCK inhibitor Y-27632 (Selleck chemicals) or StemPro-34 medium (Gibco) supplemented with 10 μM ROCK inhibitor Y-27632 (Tocris Bioscience) and 1 ng/mL rhBMP4 (R&D Systems) for 18h on an orbital shaker.

#### Day 1 (mesoderm induction)

EBs were transferred to StemPro-34 medium supplemented with 10 – 16 ng/mL rhBMP4, 6 – 10 ng/mL rhActivinA (R&D Systems;), 1 – 1.5 μM CHIR99021 (Tocris Bioscience), and 5 ng/mL rhbFGF (R&D Systems). As optimal concentrations of BMP4, ActivinA and CHIR99021 for cardiomyocyte differentiation depends on reagent lot and cell line, the indicated range of concentrations was used to achieve optimal differentiation efficiency for each cell line. *Day 3 (cardiac mesoderm specification):* EBs were harvested, washed with IMDM (Gibco), and transferred to StemPro-34 medium supplemented with 1 μM IWP2 (Tocris Bioscience) and 10 ng/mL rhVEGF (R&D Systems). *Day 6:* EBs were transferred to StemPro-34 medium supplemented with 5 ng/mL rhVEGF and cultured until day 12 with routine media changes every 3 days. *Day 12:* EBs were transferred to StemPro-34 medium without additional cytokines or RPMI-1640 medium (Gibco) supplemented with B27 (Gibco) and 2 mM L-glutamine (Gibco) and cultured until day 20 with routine media changes every 3 days. EBs were cultured at 37°C in a standard 5% CO2 incubator at 5% oxygen from day 0 – 12, and at ambient oxygen from day 12 – 20. In addition to other listed supplements, StemPro-34 medium used for differentiation was always supplemented with 50 μg/mL ascorbic acid (Sigma), 2 mM L-glutamine, 150 μg/mL transferrin (Roche, 10652202001), and 50 μg/mL monothioglycerol (Sigma). For flow cytometric analysis, hPSCs harvested at day 0 and EBs harvested at day 3 were dissociated by incubation with TrypLE for 3 – 5 min at 37°C. EBs harvested at day 20 were dissociated with 0.5 mg/mL Type II Collagenase (Worthington) in HANK’s buffer for 18 – 24h at room temperature. For measurement of transgene and cell surface marker expression, cells were stained live in FACS buffer (PBS + 5% FCS + 0.02% NaN_3_) for 15 – 30 minutes and analyzed live. For measurement of intracellular gene expression, cells were fixed with 4% PFA in PBS for 10 min at room temperature, permeabilized with 90% methanol for 20 min at 4°C, washed with PBS + 0.5% BSA and stained with unconjugated primary antibodies in FACS buffer for 18 – 24h at 4°C. Cells were washed with PBS + 0.5% BSA before staining with secondary antibodies for 30 – 60 min in FACS buffer at 4°C, washed again with PBS + 0.5% BSA and then analyzed. Samples were analyzed using an LSRFortessa (BD) or CytoFlex S (Beckman-Coulter) flow cytometer, and data analysis was done using FlowJo (TreeStar). The following antibodies were used: anti-PDGFRα-BV421 (BD, 1:25), anti-CD56-APC (BD, 1:100), anti-cTnT (ThermoFisher, 1:2000) and goat-anti-mouse-APC (BD, 1:500).

### Differentiation of hiPSCs to Myogenic Progenitors and Skeletal Myotube

hiPSCs were differentiated to the myogenic lineage to produce skeletal muscle myogenic progenitors by following the detailed protocol established by Xi *et. al.*^35^. Minor modifications to some of the reagents used are detailed here: 2% Geltrex rather than Matrigel, addition to 10 ug/ml DNase I (Millipore Sigma; 260913) to collagenase IV (ThermoFisher Scientific; 17104019) and TrypLE during the day 29 dissociation step, and SK Max medium (Wisent Bioproducts, Saint-Jean Baptise, QC, CA; 301-061-CL) as an alternative to SkGM2. Upon establishing a myogenic progenitor line, a portion of passage 0 cells were cryopreserved for future studies. To generate multinucleated myotube cultures, iPS11, NR2, and NrG derived myogenic progenitors were seeded on Geltrex coated (1:100) plates in Wisent SK Max medium supplemented with 10 % FBS and 20 ng/mL FGF2. Upon reaching 70-80% confluence the media was switched to a differentiation inducing medium consisting of DMEM with 2 % horse serum and bovine insulin (10 ug/mL) for 4-6 days. The differentiation media was exchanged every two days. Myotube cultures were fixed, stained, and imaged exactly as previously described^97^.

### Data analysis

Data analysis was conducted using both Prism (Dotmatics, Boston, MA, USA) and R. R packages used include dplyr, data.table, ggthemr, readxl, ggpubr, tidyr,stringr, ggplot2, Biostrings, Bioconductor, e1071, pbapply, stringi, stringr, rstatix, dineq, ggh4x, ggbreak and ggpattern^98^. All flow-cytometric analysis was conducted with Flowjo (BD Biosciences).

## Acknowledgements

R.A. was supported by a postdoctoral fellowship award from the Ted Rogers Centre for Heart Research. R.S. was supported by post-doctoral fellowships from the Juvenile Diabetes Research Foundation (1-PDF-2019-716 A-N) and (3-PDF-2020-954 A-N).

## Supplementary Figures

**Supplementary Figure 1:**
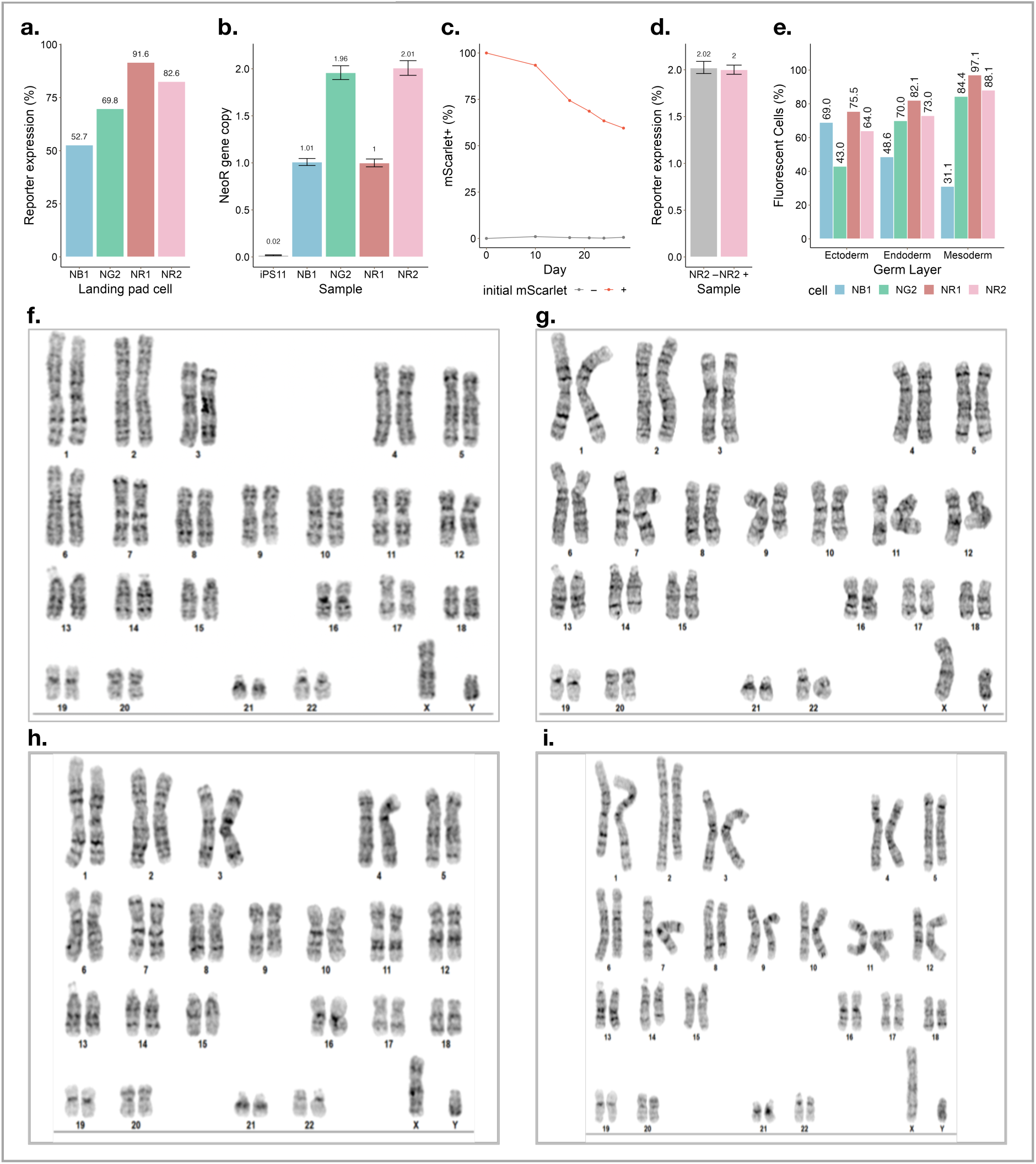
FAST-STEM hiPSC generation and profiling in pluripotent and differentiated state. **(a)** Reporter expression of tagBFP in NB1, EGFP in NG2 and mScarlet in NR1 and NR2 cells **(b)** Copy-number of FAST-STEM constructs in the human genome. **(c)** Percentage of mScarlet+ NR2 cells both initially sorted for mScarlet– and mScarlet+ sub-populations over a 28-day timecourse and **(d)** copy-number (ddPCR) of the neomycin resistance gene. **(e)** Fluorescent reporter expression post differentiation into three germ layers. Representative G-banding for **(f)** NG2 **(g)** NR2 **(h)** NB1 **(i)** NR1, where all FAST-STEM hiPSCs maintain a 46X,Y karyotype.

**Supplementary Figure 2:**
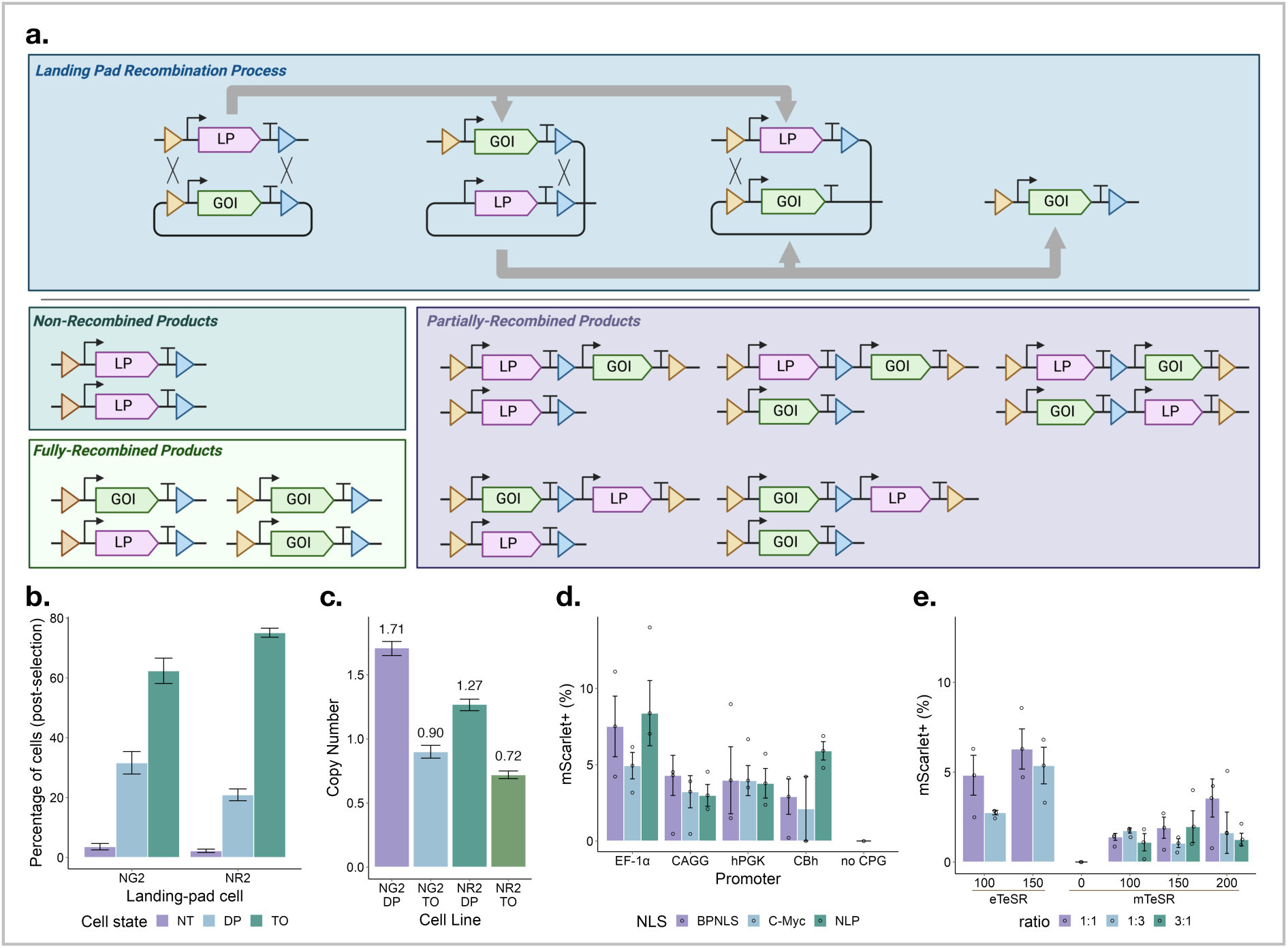
Initial results of FAST-STEM hiPSC recombination. **(a)** Schematic of transgene integration process and potential results of recombination. **(b)** Functional screening of FAST-STEM integration efficiency. Ef1a-mCherry and Ef1a-EGFP transgenes were delivered to NG2 and NR2 FAST-STEM hiPSCs respectively followed by 7-days of puromycin selection (0.25 ug/mL). NT = no transgene, DP = double-positive, TO = transgene only. **(c)** Copy-number analysis of Neomycin gene in FAST-STEM cells sorted for either double-positive or transgene-only cells. **(d)** Quantification of residual mScarlet fluorescence of transgene-delivered (EGFP+) cell populations, indicating incomplete RMCE. Combinations of four promoters (EF-1α, CAGG, hPGK, CBh) with three nuclear localization signals (NLP, C-Myc, SV40 BPNLS) were tested. **(e)** Quantification of residual mScarlet fluorescence of transgene-delivered (EGFP+) cell populations across varying transfection ratios and expansion medias, x-axis represents ng of p53DD-expression vector included in transfection mixture (of the 500 ng total DNA mixture transfected into a 24-well plate).

**Supplementary Figure 3:**
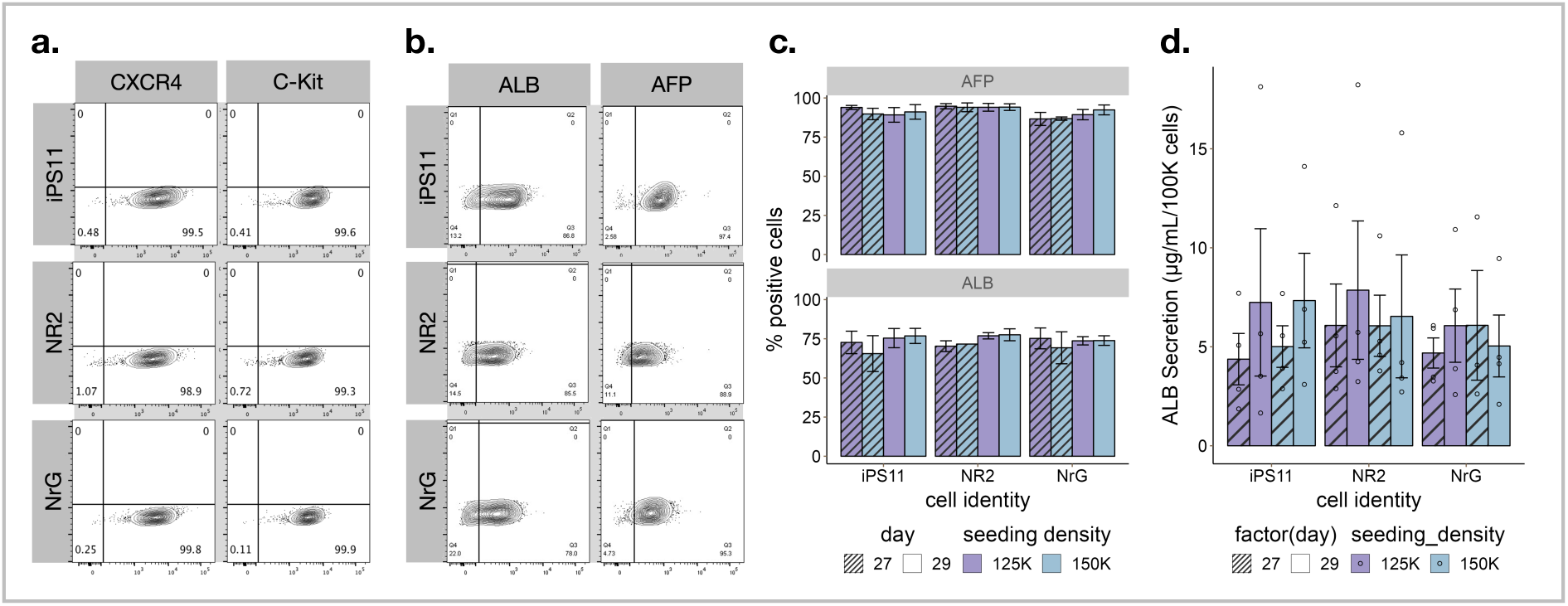
Differentiation of FAST-STEM hiPSCs into clinically relevant lineages. **(a)** Representative flow cytometry analysis of CXCR4+/CKIT+ endoderm populations differentiated from IPS11, NR5, and NRGC1 cell lines. **(b)** Quantification of the percentage of ALB+ and AFP+ hepatoblast populations at day 27 and day 29 of differentiation with different seeding densities. **(c)** Representative flow cytometry analysis of AFP+/ALB+ hepatoblast population at the end of differentiation. (**d)** Albumin levels secreted by hepatoblasts at the end of differentiation. The secretion level was detected by ELISA assay.

## REFERENCES

1. Laflamme, M. A. et al. Cardiomyocytes derived from human embryonic stem cells in pro-survival factors enhance function of infarcted rat hearts. Nat Biotechnol 25, 1015– 1024 (2007).

2. Basile, G. et al. Emerging diabetes therapies: Bringing back the β-cells. Mol Metab 60, 101477 (2022).

3. BlueRock Therapeutics. Phase 1 Safety and Tolerability Study of MSK-DA01 Cell Therapy for Advanced Parkinson’s Disease. https://clinicaltrials.gov/study/NCT04802733 Preprint at (2022).

4. Leone, M. A. et al. Phase I clinical trial of intracerebroventricular transplantation of allogeneic neural stem cells in people with progressive multiple sclerosis. Cell Stem Cell 30, 1597–1609.e8 (2023).

5. Morsut, L. et al. Engineering Customized Cell Sensing and Response Behaviors Using Synthetic Notch Receptors. Cell 164, 780–791 (2016).

6. Amos, M. & Goñi-Moreno, A. Cellular Computing and Synthetic Biology BT - Computational Matter. in (eds. Stepney, S., Rasmussen, S. & Amos, M.) 93–110 (Springer International Publishing, Cham, 2018). doi:10.1007/978-3-319-65826-1_7.

7. Schwarz, K. A., Daringer, N. M., Dolberg, T. B. & Leonard, J. N. Rewiring human cellular input–output using modular extracellular sensors. Nat Chem Biol 13, 202–209 (2017).

8. Mansouri, M., Strittmatter, T. & Fussenegger, M. Light-Controlled Mammalian Cells and Their Therapeutic Applications in Synthetic Biology. Advanced Science 6, 1800952 (2019).

9. Mansouri, M. & Fussenegger, M. Electrogenetics: Bridging synthetic biology and electronics to remotely control the behavior of mammalian designer cells. Curr Opin Chem Biol 68, 102151 (2022).

10. Nims, R. J., Pferdehirt, L. & Guilak, F. Mechanogenetics: harnessing mechanobiology for cellular engineering. Curr Opin Biotechnol 73, 374–379 (2022).

11. Dhahri, W. et al. In Vitro Matured Human Pluripotent Stem Cell–Derived Cardiomyocytes Form Grafts With Enhanced Structure and Function in Injured Hearts. Circulation 145, 1412–1426 (2022).

12. Choi, J. et al. A time-resolved, multi-symbol molecular recorder via sequential genome editing. Nature 608, 98–107 (2022).

13. Wang, N. B., Beitz, A. M. & Galloway, K. Engineering cell fate: Applying synthetic biology to cellular reprogramming. Curr Opin Syst Biol 24, 18–31 (2020).

14. Mansouri, M., Ray, P. G., Franko, N., Xue, S. & Fussenegger, M. Design of programmable post-translational switch control platform for on-demand protein secretion in mammalian cells. Nucleic Acids Res 51, e1–e1 (2023).

15. Park, J. S. et al. Synthetic control of mammalian-cell motility by engineering chemotaxis to an orthogonal bioinert chemical signal. Proc Natl Acad Sci U S A 111, 5896–5901 (2014).

16. Scheller, L. & Fussenegger, M. From synthetic biology to human therapy: engineered mammalian cells. Curr Opin Biotechnol 58, 108–116 (2019).

17. Lonzarić, J., Fink, T. & Jerala, R. Design and applications of synthetic information processing circuits in mammalian cells. in Synthetic Biology: Volume 2 vol. 2 1–34 (The Royal Society of Chemistry, 2018).

18. Mukherji, S. & van Oudenaarden, A. Synthetic biology: understanding biological design from synthetic circuits. Nat Rev Genet 10, 859–871 (2009).

19. Mansouri, M. & Fussenegger, M. Therapeutic cell engineering: designing programmable synthetic genetic circuits in mammalian cells. Protein Cell 13, 476–489 (2022).

20. Linder, J., Srivastava, D., Yuan, H., Agarwal, V. & Kelley, D. R. Predicting RNA-seq coverage from DNA sequence as a unifying model of gene regulation. bioRxiv 2023.08.30.555582 (2023) doi:10.1101/2023.08.30.555582.

21. Zrimec, J. et al. Controlling gene expression with deep generative design of regulatory DNA. Nat Commun 13, 5099 (2022).

22. Lyu, S., Sowlati-Hashjin, S. & Garton, M. Variational autoencoder for design of synthetic viral vector serotypes. Nat Mach Intell 6, 147–160 (2024).

23. Ichikawa, D. M. et al. A universal deep-learning model for zinc finger design enables transcription factor reprogramming. Nat Biotechnol 41, 1117–1129 (2023).

24. Capponi, S. & Daniels, K. G. Harnessing the power of artificial intelligence to advance cell therapy. Immunol Rev 320, 147–165 (2023).

25. Wang, H. et al. GPro: generative AI-empowered toolkit for promoter design. Bioinformatics 40, btae123 (2024).

26. Repecka, D. et al. Expanding functional protein sequence spaces using generative adversarial networks. Nat Mach Intell 3, 324–333 (2021).

27. Nguyen, E. et al. Sequence modeling and design from molecular to genome scale with Evo. bioRxiv 2024.02.27.582234 (2024) doi:10.1101/2024.02.27.582234.

28. de Almeida, B. P., et al. SegmentNT: annotating the genome at single-nucleotide resolution with DNA foundation models. bioRxiv 2024.03.14.584712 (2024) doi:10.1101/2024.03.14.584712.

29. DaSilva, L. F. et al. DNA-Diffusion: Leveraging Generative Models for Controlling Chromatin Accessibility and Gene Expression via Synthetic Regulatory Elements. bioRxiv 2024.02.01.578352 (2024) doi:10.1101/2024.02.01.578352.

30. Di Blasi, R. et al. Resource-aware construct design in mammalian cells. Nat Commun 14, 3576 (2023).

31. Baker, M., Reynolds, H. M., Lumicisi, B. & Bryson, C. J. Immunogenicity of protein therapeutics: The key causes, consequences and challenges. Self Nonself 1, 314–322 (2010).

32. Cabrera, A. et al. The sound of silence: Transgene silencing in mammalian cell engineering. Cell Syst 13, 950–973 (2022).

33. Shakiba, N., Jones, R. D., Weiss, R. & Del Vecchio, D. Context-aware synthetic biology by controller design: Engineering the mammalian cell. Cell Syst 12, 561–592 (2021).

34. Hulme, A. J., Maksour, S., St-Clair Glover, M., Miellet, S. & Dottori, M. Making neurons, made easy: The use of Neurogenin-2 in neuronal differentiation. Stem Cell Reports 17, 14–34 (2022).

35. Xi, H. et al. In-Vivo Human Somitogenesis Guides Somite Development from hPSCs. Cell Rep 18, 1573–1585 (2017).

36. Zare, M. et al. Efficient biotechnological approach for lentiviral transduction of induced pluripotent stem cells. Artif Cells Nanomed Biotechnol 44, 743–748 (2016).

37. Park, M. A., Jung, H. S. & Slukvin, I. Genetic Engineering of Human Pluripotent Stem Cells Using PiggyBac Transposon System. Curr Protoc Stem Cell Biol 47, e63–e63 (2018).

38. Hoffmann, D. et al. Detailed comparison of retroviral vectors and promoter configurations for stable and high transgene expression in human induced pluripotent stem cells. Gene Ther 24, 298–307 (2017).

39. Ellis, J. Silencing and variegation of gammaretrovirus and lentivirus vectors. Hum Gene Ther 16, 1241–1246 (2005).

40. Park, I.-H. et al. Disease-specific induced pluripotent stem cells. Cell 134, 877–886 (2008).

41. Mali, P. et al. RNA-guided human genome engineering via Cas9. Science 339, 823– 826 (2013).

42. Ihry, R. J. et al. p53 inhibits CRISPR-Cas9 engineering in human pluripotent stem cells. Nat Med 24, 939–946 (2018).

43. Chi, X., Zheng, Q., Jiang, R., Chen-Tsai, R. Y. & Kong, L.-J. A system for site-specific integration of transgenes in mammalian cells. PLoS One 14, e0219842 (2019).

44. Ran, B. et al. A versatile platform for locus-scale genome rewriting and verification. Proceedings of the National Academy of Sciences 118, e2023952118 (2021).

45. Zhu, F. et al. DICE, an efficient system for iterative genomic editing in human pluripotent stem cells. Nucleic Acids Res 42, e34–e34 (2014).

46. Matreyek, K. A., Stephany, J. J., Chiasson, M. A., Hasle, N. & Fowler, D. M. An improved platform for functional assessment of large protein libraries in mammalian cells. Nucleic Acids Res 48, e1–e1 (2020).

47. Gaidukov, L. et al. A multi-landing pad DNA integration platform for mammalian cell engineering. Nucleic Acids Res 46, 4072–4086 (2018).

48. Inniss, M. C. et al. A novel Bxb1 integrase RMCE system for high fidelity site-specific integration of mAb expression cassette in CHO Cells. Biotechnol Bioeng 114, 1837– 1846 (2017).

49. Cerbini, T. et al. Transcription activator-like effector nuclease (TALEN)-mediated CLYBL targeting enables enhanced transgene expression and one-step generation of dual reporter human induced pluripotent stem cell (iPSC) and neural stem cell (NSC) lines. PLoS One 10, e0116032–e0116032 (2015).

50. Pellenz, S. et al. New Human Chromosomal Sites with ‘Safe Harbor’ Potential for Targeted Transgene Insertion. Hum Gene Ther 30, 814–828 (2019).

51. Irion, S. et al. Identification and targeting of the ROSA26 locus in human embryonic stem cells. Nat Biotechnol 25, 1477–1482 (2007).

52. Kim, J. et al. Cre/Lox-based RMCE for Site-specific Integration in CHO Cells. Biotechnology and bioprocess engineering 26, 795–803 (2021).

53. Leonard, S. P. et al. Genetic Engineering of Bee Gut Microbiome Bacteria with a Toolkit for Modular Assembly of Broad-Host-Range Plasmids. ACS Synth Biol 7, 1279–1290 (2018).

54. Lee, M. E., DeLoache, W. C., Cervantes, B. & Dueber, J. E. A Highly Characterized Yeast Toolkit for Modular, Multipart Assembly. ACS Synth Biol 4, 975–986 (2015).

55. Weber, E., Engler, C., Gruetzner, R., Werner, S. & Marillonnet, S. A Modular Cloning System for Standardized Assembly of Multigene Constructs. PLoS One 6, e16765- (2011).

56. Fonseca, J. P. et al. A Toolkit for Rapid Modular Construction of Biological Circuits in Mammalian Cells. ACS Synth Biol 8, 2593–2606 (2019).

57. Qin, J. Y. et al. Systematic Comparison of Constitutive Promoters and the Doxycycline-Inducible Promoter. PLoS One 5, e10611- (2010).

58. Luo, Y. et al. Stable enhanced green fluorescent protein expression after differentiation and transplantation of reporter human induced pluripotent stem cells generated by AAVS1 transcription activator-like effector nucleases. Stem Cells Transl Med 3, 821–835 (2014).

59. Norrman, K. et al. Quantitative Comparison of Constitutive Promoters in Human ES cells. PLoS One 5, e12413- (2010).

60. Liao, W. et al. Functional Characterization of Insulation Effect for Synthetic Gene Circuits in Mammalian Cells. ACS Synth Biol 7, 412–418 (2018).

61. Fonseca, J. P. et al. A Toolkit for Rapid Modular Construction of Biological Circuits in Mammalian Cells. ACS Synth Biol 8, 2593–2606 (2019).

62. Li, M. et al. Transient inhibition of p53 enhances prime editing and cytosine base-editing efficiencies in human pluripotent stem cells. Nat Commun 13, 6354 (2022).

63. Hong, H. et al. Suppression of induced pluripotent stem cell generation by the p53–p21 pathway. Nature 460, 1132–1135 (2009).

64. Anzalone, A. V. et al. Search-and-replace genome editing without double-strand breaks or donor DNA. Nature 576, 149–157 (2019).

65. Li, W. et al. Quality control, modeling, and visualization of CRISPR screens with MAGeCK-VISPR. Genome Biol 16, 281 (2015).

66. Ogawa, S. et al. Three-dimensional culture and cAMP signaling promote the maturation of human pluripotent stem cell-derived hepatocytes. Development 140, 3285–3296 (2013).

67. Yung, T. et al. Sufu- and Spop-mediated downregulation of Hedgehog signaling promotes beta cell differentiation through organ-specific niche signals. Nat Commun 10, 4647 (2019).

68. Hogrebe, N. J., Augsornworawat, P., Maxwell, K. G., Velazco-Cruz, L. & Millman, J. R. Targeting the cytoskeleton to direct pancreatic differentiation of human pluripotent stem cells. Nat Biotechnol 38, 460–470 (2020).

69. Balboa, D. et al. Functional, metabolic and transcriptional maturation of human pancreatic islets derived from stem cells. Nat Biotechnol 40, 1042–1055 (2022).

70. Kattman, S. J. et al. Stage-Specific Optimization of Activin/Nodal and BMP Signaling Promotes Cardiac Differentiation of Mouse and Human Pluripotent Stem Cell Lines. Cell Stem Cell 8, 228–240 (2011).

71. Lee, J. H., Protze, S. I., Laksman, Z., Backx, P. H. & Keller, G. M. Human Pluripotent Stem Cell-Derived Atrial and Ventricular Cardiomyocytes Develop from Distinct Mesoderm Populations. Cell Stem Cell 21, 179–194.e4 (2017).

72. Xu, Z. et al. Accuracy and efficiency define Bxb1 integrase as the best of fifteen candidate serine recombinases for the integration of DNA into the human genome. BMC Biotechnol 13, 87 (2013).

73. Blanch-Asensio, A. et al. Generation of AAVS1 and CLYBL STRAIGHT-IN v2 acceptor human iPSC lines for integrating DNA payloads. Stem Cell Res 66, 102991 (2023).

74. Liu, J., Jeppesen, I., Nielsen, K. & Jensen, T. G. φc31 integrase induces chromosomal aberrations in primary human fibroblasts. Gene Ther 13, 1188–1190 (2006).

75. Ehrhardt, A., Engler, J. A., Xu, H., Cherry, A. M. & Kay, M. A. Molecular Analysis of Chromosomal Rearrangements in Mammalian Cells After øC31-Mediated Integration. Hum Gene Ther 17, 1077–1094 (2006).

76. Hockemeyer, D. et al. Efficient targeting of expressed and silent genes in human ESCs and iPSCs using zinc-finger nucleases. Nat Biotechnol 27, 851–857 (2009).

77. Ranawakage, D. C. et al. Efficient CRISPR-Cas9-Mediated Knock-In of Composite Tags in Zebrafish Using Long ssDNA as a Donor. Front Cell Dev Biol 8, (2021).

78. Li, X.-L. et al. Highly efficient genome editing via CRISPR–Cas9 in human pluripotent stem cells is achieved by transient BCL-XL overexpression. Nucleic Acids Res 46, 10195–10215 (2018).

79. Haideri, T., et al. Robust genome editing via modRNA-based Cas9 or base editor in human pluripotent stem cells. Cell Reports Methods 2, 100290 (2022).

80. Gaidukov, L. et al. A multi-landing pad DNA integration platform for mammalian cell engineering. Nucleic Acids Res 46, 4072–4086 (2018).

81. Roelle, S. M., Kamath, N. D. & Matreyek, K. A. Mammalian Genomic Manipulation with Orthogonal Bxb1 DNA Recombinase Sites for the Functional Characterization of Protein Variants. ACS Synth Biol 12, 3352–3365 (2023).

82. Hart, T. et al. Evaluation and Design of Genome-Wide CRISPR/SpCas9 Knockout Screens. G3 Genes|Genomes|Genetics 7, 2719–2727 (2017).

83. Sack, L. M., Davoli, T., Xu, Q., Li, M. Z. & Elledge, S. J. Sources of Error in Mammalian Genetic Screens. G3 Genes|Genomes|Genetics 6, 2781–2790 (2016).

84. Xie, S., Cooley, A., Armendariz, D., Zhou, P. & Hon, G. C. Frequent sgRNA-barcode recombination in single-cell perturbation assays. PLoS One 13, e0198635- (2018).

85. Hill, A. J. et al. On the design of CRISPR-based single-cell molecular screens. Nat Methods 15, 271–274 (2018).

86. Xiong, K. et al. An optimized genome-wide, virus-free CRISPR screen for mammalian cells. Cell Reports Methods 1, 100062 (2021).

87. Matreyek, K. A., Stephany, J. J. & Fowler, D. M. A platform for functional assessment of large variant libraries in mammalian cells. Nucleic Acids Res 45, e102–e102 (2017).

88. Gibson, D. G. et al. Enzymatic assembly of DNA molecules up to several hundred kilobases. Nat Methods 6, 343–345 (2009).

89. Tian, S., Yesselman, J. D., Cordero, P. & Das, R. Primerize: automated primer assembly for transcribing non-coding RNA domains. Nucleic Acids Res 43, W522– W526 (2015).

90. Tian, S. & Das, R. Primerize-2D: automated primer design for RNA multidimensional chemical mapping. Bioinformatics 33, 1405–1406 (2017).

91. Li, S., Prasanna, X., Salo, V. T., Vattulainen, I. & Ikonen, E. An efficient auxin-inducible degron system with low basal degradation in human cells. Nat Methods 16, 866–869 (2019).

92. Ran, F. A. et al. Genome engineering using the CRISPR-Cas9 system. Nat Protoc 8, 2281–2308 (2013).

93. Rosenstein, A. H. & Walker, V. K. Fidelity of a Bacterial DNA Polymerase in Microgravity, a Model for Human Health in Space. Front Cell Dev Biol 9, (2021).

94. Nostro, M. C. et al. Efficient Generation of NKX6-1+ Pancreatic Progenitors from Multiple Human Pluripotent Stem Cell Lines. Stem Cell Reports 4, 591–604 (2015).

95. Dubois, N. C. et al. SIRPA is a specific cell-surface marker for isolating cardiomyocytes derived from human pluripotent stem cells. Nat Biotechnol 29, 1011– 1018 (2011).

96. Kawada, J. et al. Generation of a Motor Nerve Organoid with Human Stem Cell-Derived Neurons. Stem Cell Reports 9, 1441–1449 (2017).

97. Ebrahimi, M. et al. De novo revertant fiber formation and therapy testing in a 3D culture model of Duchenne muscular dystrophy skeletal muscle. Acta Biomater 132, 227–244 (2021).

98. Xu, S. et al. Use ggbreak to Effectively Utilize Plotting Space to Deal With Large Datasets and Outliers. Front Genet 12, (2021).

